# Temperature Effect on Polymerase Fidelity

**DOI:** 10.1101/2020.08.04.236919

**Authors:** Yuan Xue, Ido Braslavsky, Stephen R. Quake

## Abstract

The discovery of extremophiles helped enable the development of groundbreaking technology such as polymerase chain reaction. Temperature variation is often an essential step of these technology platforms, but the effect of temperature on the error rate of polymerases from different origins is under-explored. Here, we applied high-throughput sequencing to profile the error rates of DNA polymerases from psychrophilic, mesophilic, and thermophilic origins with single-molecule resolution. We found that reaction temperature substantially increases substitution and deletion error rates of psychrophilic and mesophilic DNA polymerases. Our motif analysis shows that the substitution error profiles cluster according to phylogenetic similarity of polymerases, not reaction temperature, thus suggesting that reaction temperature increases global error rate of polymerases independent of sequence context. Intriguingly, we also found that the DNA polymerase I of a psychrophilic bacteria exhibits higher polymerization activity than its mesophilic ortholog across all temperature ranges, including down to −19°C which is well below the freezing temperature of water. Our results provide a useful reference for how reaction temperature, a crucial parameter of biochemistry, can affect DNA polymerase fidelity in organisms adapted to a wide range of thermal environments.

## Introduction

Micro-organisms which live in cold environment are confronted with the thermodynamic challenge of maintaining the chemical processes of life; however, the abundance of various life-forms found in near or below freezing temperature suggests that biology has evolved ways to overcome such obstacles. Contrary to what one would expect from naïve application of the Arrhenius equation, micro-organisms in cold environments do not grow exponentially slower in the cold than their mesophilic cousins at room temperature ^1,2^. This suggests that psychrophilic enzymes have evolved to catalyze relevant activities at low temperature ^3–5^. Low-temperature adaptation is thought to have increased the catalytic rate of enzymes by increasing local flexibility of their active sites, thus lowering activation energy barriers ^6–9^. This can be contrasted with high-temperature adaptation which confers structural robustness to thermal denaturation by increasing intra-molecular interactions that maintain an enzyme’s functional tertiary shape. Studies based on directed evolution have shown that structural stability and enzymatic efficiency can be optimized simultaneously and are thought to be mutually compatible properties ^10,11^. Combining these observations helps explain the general biochemical trend of cold-adapted enzymes: an increase in enzyme’s catalytic activity is often associated with a reduced binding affinity and a broader substrate specificity ^12,13^. However, a direct test of these ideas on orthologous enzymes from organisms adapted to low, medium, and high temperatures has yet been performed.

As DNA polymerase is a central component of cellular replication and biotechnological platforms ^14^, understanding how reaction temperature can affect fidelity is vital. Polymerase error rate is tightly controlled to ensure the successful duplication of genetic material. Replication error from DNA polymerase is a source of genetic variations and underlies the cause of many diseases ^15–18^. Polymerase errors also play a crucial role in biotechnology applications such as polymerase chain reaction (PCR) and multiple-displacement amplification (MDA) ^19–21^. Polymerase can introduce errors at an exponential rate, significantly impairing downstream interpretation. In high-throughput sequencing, polymerase replication errors can reduce base calling accuracy and precision ^22–24^. The increasing applications of nucleic acid amplification underscore a need to better understand the degree to which physical parameters, such as temperature, affects polymerase fidelity.

This study set out to provide a reference for how temperature affects the activity and error profiles of DNA polymerases of psychrophilic, mesophilic, and thermophilic origins (**Figure 1a**). Previous studies reported psychrophilic DNA polymerases’ activity at ambient temperature ^25,26^; however, the behavior of these polymerases across a wide range of reaction temperatures remains unexplored. In this study, we demonstrate for the first time that DNA polymerase I from a psychrophilic bacteria, *Psychromonas ingrahamii*, retains DNA replication activity below water’s freezing temperature. We used random nucleotides to multiplex single-molecule measurement of insertion, deletion, and substitution events for DNA polymerases from different origins across a range of temperature conditions. This enabled a comprehensive mapping of DNA polymerase error profiles as a function of reaction temperature, revealing an unexpected relationship between fidelity and temperature adaptation. Our report provides an important reference to inform design choices for future biotechnological applications of DNA polymerase.

**Figure 1.**
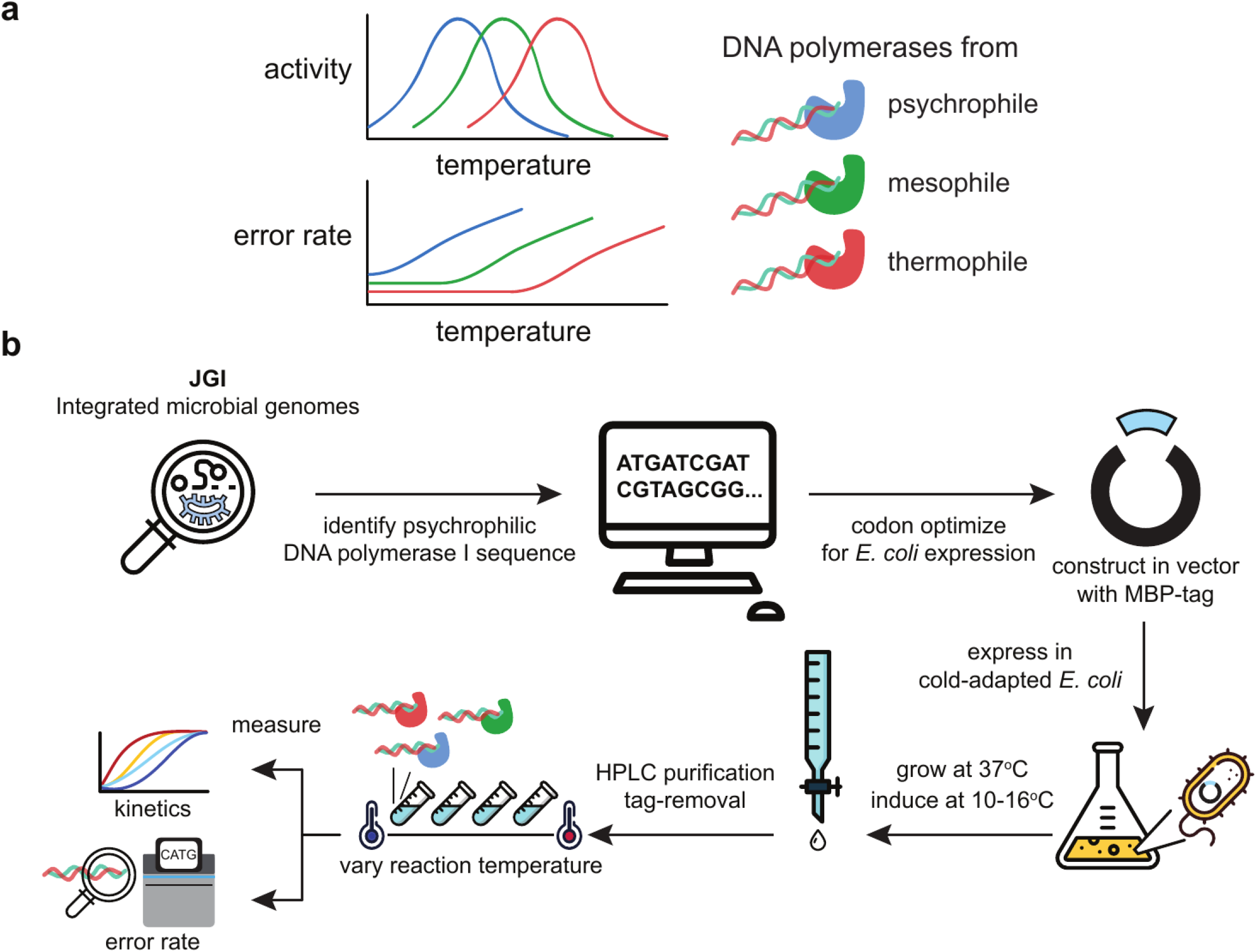
**(a)** A schematic illustrating the apparent trade-off between enzymatic speed versus fidelity in the context of temperature adaptation. Thermophilic enzymes are expected to be selected for low error rate due to increased environmental temperature. Psychrophilic enzymes are expected to be selected for high enzymatic speed due to decreased environmental temperature. **(b)** Graphical summary of the study.

## Results

### Psychrophilic DNA polymerase is more active than mesophilic and thermophilic polymerases and can replicate DNA below zero degrees Celsius

To address whether there is an activity and fidelity trade-off in DNA polymerase, we wanted to measure and compare the temperature-dependence of its activity from a wide range of microbial sources. As psychrophilic DNA polymerase is unavailable commercially, we set out to recombinantly purify one for our study. Based on homology search, we identified DNA polymerase I (PIPI; IMG Gene ID: 639798289), a family A polymerase, from *Psychromonas ingrahamii*,a gram-negative bacteria that can grow and replicate at −12°C in laboratory condition ^31–33^. MBP-PIPI was purified by maltose affinity column, cleaved off its tandem-fused MBP tag, and then re-purified with size exclusion chromatography (**Methods**) (**Supplemental Figure 1a**). A graphical overview of our study is summarized in **Figure 1b**.

To determine the temperature profile of polymerase extension activity, we measured the extension rate of DNA polymerase I of psychrophilic, mesophilic, and thermophilic origins across a range of incubation temperatures. Briefly, we use a fluorometric assay to monitor the level of fluorescence over time at different incubation temperature. When the polymerase extends from primed template, we can detect a increase of fluorescence emission due to binding of double-stranded DNA to the EvaGreen fluorophore. As the fluorescence level is linearly proportional to the amount of dsDNA product, we can thus infer the polymerase extension activity by calculating the initial rate of fluorescence change per unit time (**Methods**). We were able to detect robust activity in the recombinantly purified PIPI (left panel in **Figure 2a**), suggesting it is biochemically active. Similar to its mesophilic ortholog, Klenow fragment (KleLF), PIPI exhibits peak activity at around 37°C. This is consistent with previous reports that the optimal temperature of enzyme activity can differ from the temperature to which the host organism is adapted ^26,34^. Klenow fragment lacking 3′**→**5′ exonuclease domain (KleExo-) exhibits higher activity across all temperature range, suggesting that the observed activity may also be dependent on the presence of exonuclease domain. As expected, Taq DNA polymerase is inactive at low temperature below 30°C and its activity continues to increase at up to 72°C. Strikingly, PIPI exhibits higher extension activity than KleLF at or below 37°C (**Figure 2b**). The fold difference in activity between PIPI and Taq is even more dramatic, displaying over a 10-fold difference at 30°C. PIPI activity decreases with increasing reaction temperature due to thermal inhibition. Our observation is thus consistent with the hypothesis that enzymes of psychrophilic origin are evolved to have higher activity, even at moderate temperatures.

**Figure 2.**
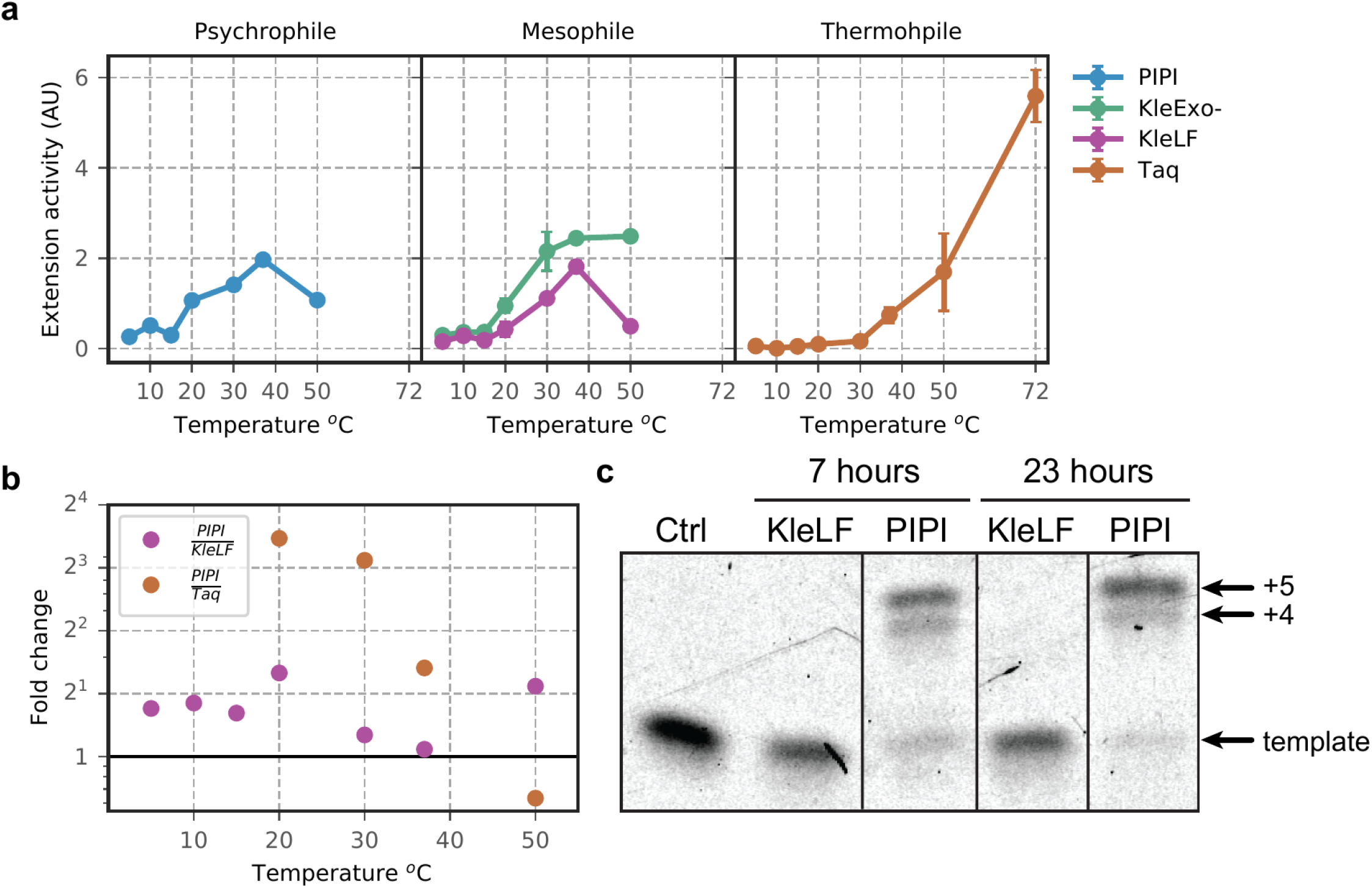
**(a)** Extension activity of purified psychrophilic, mesophilic, and thermophilic DNA polymerases. **(b)** Fold change of PIPI extension activity relative to KleLF or Taq. Taq activity is indistinguishable from background below 20°C so comparison to its activity is excluded at low temperature range. **(c)** PIPI and KleLF are pre-diluted to equal concentration and they are co-incubated with a primed template at −19°C for 7 or 23 hours. DNA template (22 bp) is hybridized with a complementary 17 bp long 5′-ATTOTM 633 labeled oligonucleotide, leaving 5 bp gap at the 3′ end.

Previous reports showed that *Psychromonas ingrahamii* could grow and replicate at negative 12°C ^32^. We thus reasoned that PIPI should remain active below 0°C. Using 5’-fluorescently labeled primers (**Methods**), we performed a gel extension assay to measure the activity of PIPI and KleLF at −19°C in an aqueous solution that had been supplemented with 30% glycerol. Strikingly, we discover that PIPI retains extension activity as it completed extension of a primed template with a 5-nucleotide gap under 7 hours, while KleLF did not (**Figure 2c**). KleLF failed to complete extension at −19°C up to 23 hours (**Figure 2c**). We wondered whether this is due to inhibition of KleLF binding to dsDNA. By modeling the thermodynamics of binding in previous data, we found that the KleLF binds to dsDNA favorably at down to −19°C (ΔG: −9.48 kcal/mol)^35^. This suggests KleLF is catalytically inactivated at low temperature, as suggested by other studies ^34,36^. In the later part of this article, we discuss several exciting possibility of utilizing PIPI for sub-zero reactions that may open the door for novel biochemical applications.

### Reaction temperature increases the error rate of DNA polymerases

In order to profile polymerase errors across a range of reaction temperatures and conditions, we adapted a high-throughput multiplexed sequencing approach ^29^. Briefly, a reaction mixture that contains a complementary primer and single-stranded DNA (ssDNA) pUC19 fragment (NEB) is equilibrated to the desired temperature prior to the addition of polymerase. Unless otherwise noted, the mixture is buffered against 3-(*N*-morpholino)propanesulfonic acid (MOPS, pH 8.5) whose pKa has relatively small temperature-dependence. It is essential to control for changes in pH as this can influence enzyme fidelity independently of temperature ^37^. After quenching, dsDNA products are PCR amplified, purified, and pooled at equimolar concentrations for sequencing on a MiSeq 2 × 300 bp platform (**Methods**). We found no runoff amplification by-products in up to 26 rounds of PCR amplification, while 20 rounds of PCR amplification failed to yield sufficient DNA amount for downstream purification (**Supplemental Figure 2a**). The purified libraries are pooled and sequenced on the Illumina MiSeq platform. We deconvolved the sequenced reads by their reaction indices and UMIs, removed sequencing errors by trimming reads with low quality, and called consensus sequence of molecules. Quantification of base counts with high sequence consensus reveals uniform coverage along the entire pUC19 template, indicating no substantial positional effect on measurement dropout. An overview of the experimental and analytical workflows is illustrated in **Figure 3a**.

**Figure 3.**
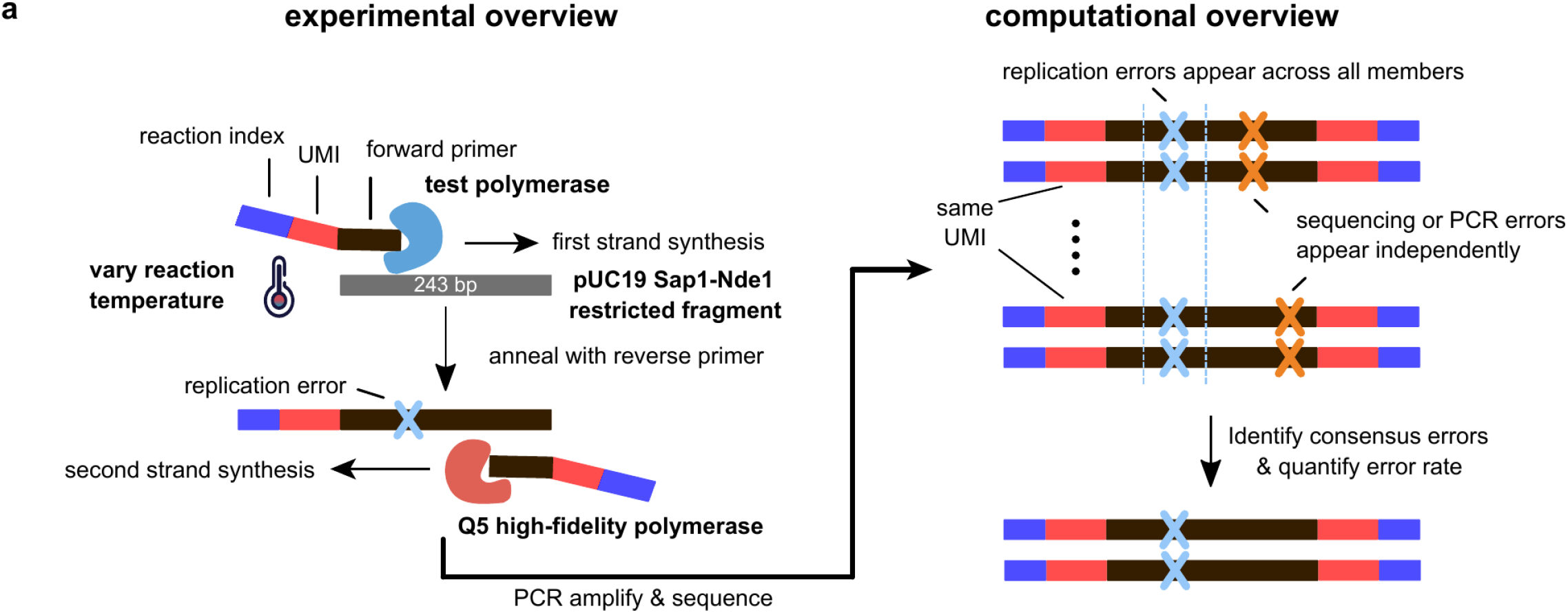
**(a)** A schematic illustrating the experimental overview of multiplexed single-molecule measurement of polymerase error rate at various reaction temperature. Replication errors (highlighted by blue cross) are distinguished from sequencing errors (highlighted by orange cross) by comparing consensus sequence of individual molecules to the reference sequence in pUC19 Sap1-Nde1 fragment.

We categorized replication error events as either substitution, deletion or insertion error and quantified the frequency of these error types in each unique molecule. As insertion and deletion events are relatively rare, we focused our analysis primarily on substitution error of psychrophilic, mesophilic, and thermophilic DNA polymerases in **Figures 4a**. A comprehensive report of error rates and coverage for thermophilic, psychrophilic, and mesophilic DNA polymerases is presented in **Tables 1 – 3**. Quantification of Q5 DNA polymerase error rate in its native buffer (Q5NativeBuffer) at 72°C provides a baseline for our measurement, revealing an average substitution, deletion, and insertion rates of 4.44 × 10^−6^, 9.50 × 10^−7^, and 1.37 × 10^−7^ per base, respectively, which agrees well with previously reported values (**Table 1**) ^29,38^. Phusion (ThermoFisher), another high-fidelity and commonly used thermophilic polymerase, has a similar substitution rate of 5.05 × 10^−6^ per base (**Table 1**) at 72°C in its native Phusion HF buffer (PhusionNativeBuffer; ThermoFisher). We discover that the substitution rate of Phusion DNA polymerase increases by more than a factor of two when switched from its native buffer to MOPS; furthermore, we notice an increased substitution rate of Phusion polymerase in its native buffer (PhusionNativeBuffer) in lower reaction temperature. We suspect the native buffer of Phusion polymerase (ThermoFisher) is designed for a specific temperature and its pH is highly temperature-dependent. In low temperatures, the pH changes in the buffer system causes sub-optimal polymerase fidelity. This would explain why the substitution rate of Phusion in the MOPS buffer remains relatively invariant to temperature changes (**Table 1** and right panel in **Figure 4a**). Supporting this hypothesis, we do not observe substantial differences in the substitution rate of Q5 polymerase in MOPS buffer between 30°C and 72°C. Our finding strongly suggests that thermophilic polymerases from commercial sources should use alternative buffers to ensure optimal fidelity at low temperatures.

**Figure 4.**
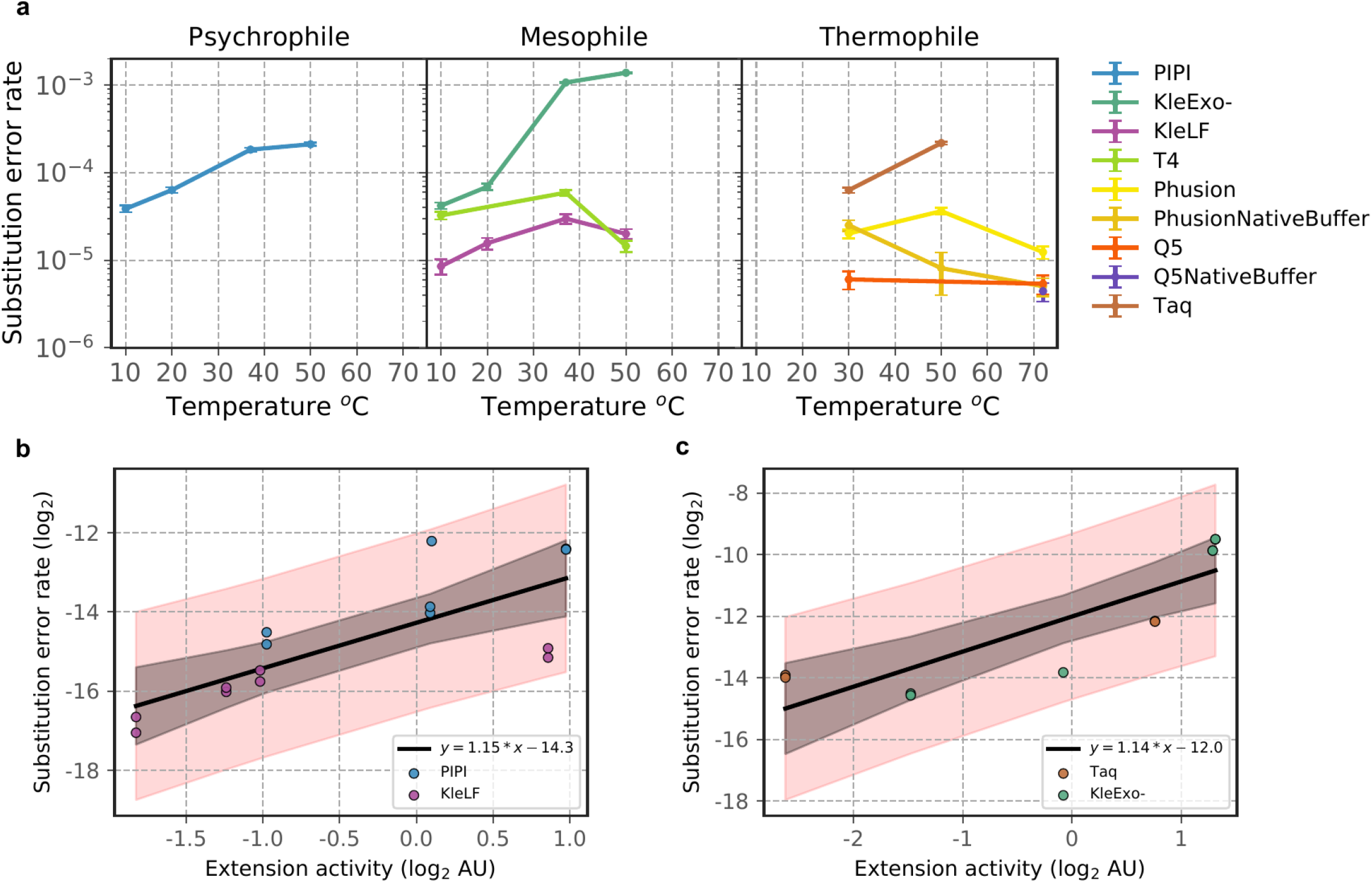
**(a)** Substitution error rates (per base) of psychrophilic (left), mesophilic (center), and thermophilic (right) DNA polymerases as a function of reaction temperature. Error bars are standard deviation of the mean error rate fitted to a Binomial distribution. Linear regression of substitution error rate on extension activity for polymerases with predicted exonuclease activity **(b)** (PIPI and KleLF) and **(c)** for ones without (Taq and KleExo-). Shaded regions indicate 95% (black) prediction intervals of linear regression and (red) confidence intervals of sampling mean.

**Table 1.** Error rates (per base) for thermophilic DNA polymerases.

| Origin | Reaction | Temperature (°C) | Substitution rate (base<sup>-1</sup>) | Deletion rate (base<sup>-1</sup>) | Insertion rate (base<sup>-1</sup>) | Bases counted |
| --- | --- | --- | --- | --- | --- | --- |
| Thermophile | Q5<sup>NB</sup> | 72 | $4.44 \times 10^{-6}$ | $9.50 \times 10^{-7}$ | $1.37 \times 10^{-7}$ | 15,536,120 |
| | Q5 | 30 | $6.04 \times 10^{-6}$ | $7.45 \times 10^{-7}$ | $5.31 \times 10^{-7}$ | 6,001,934 |
| | Q5 | 72 | $5.38 \times 10^{-6}$ | ND | ND | 6,340,445 |
| | Taq | 30 | $6.32 \times 10^{-5}$ | $2.27 \times 10^{-6}$ | $2.01 \times 10^{-6}$ | 6,497,391 |
| | Taq | 50 | $2.18 \times 10^{-4}$ | $2.18 \times 10^{-5}$ | $3.53 \times 10^{-6}$ | 6,320,921 |
| | Phusion<sup>NB</sup> | 30 | $2.51 \times 10^{-5}$ | ND | $8.32 \times 10^{-7}$ | 2,064,657 |
| | Phusion<sup>NB</sup> | 50 | $8.11 \times 10^{-6}$ | ND | ND | 478,302 |
| | Phusion<sup>NB</sup> | 72 | $5.05 \times 10^{-6}$ | ND | ND | 10,515,198 |
| | Phusion | 30 | $2.02 \times 10^{-5}$ | $2.32 \times 10^{-6}$ | $4.81 \times 10^{-7}$ | 6,479,246 |
| | Phusion | 50 | $3.64 \times 10^{-5}$ | $2.46 \times 10^{-6}$ | $4.84 \times 10^{-7}$ | 7,260,714 |
| | Phusion | 72 | $1.24 \times 10^{-5}$ | ND | $5.11 \times 10^{-7}$ | 6,182,116 |

<sup>NB</sup>=reaction carried out in native buffer supplied by the manufacturer.
<sup>ND</sup>=error frequency below the detectable error baseline.

**Table 2.**
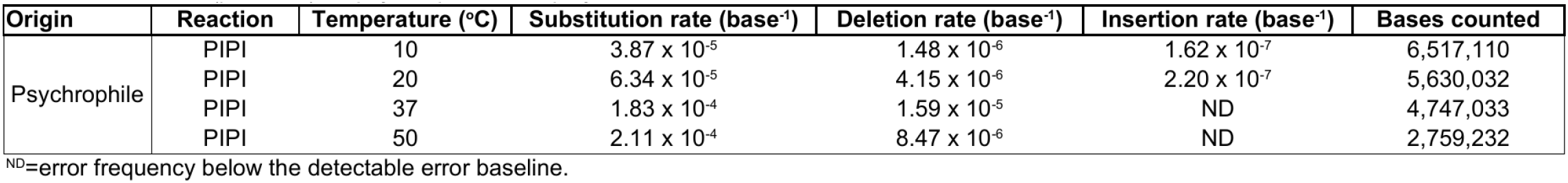
Error rates (per base) for psychrophilic DNA polymerase.

**Table 3.** Error rates (per base) for mesophilic DNA polymerases.

| Origin | Reaction | Temperature (°C) | Substitution rate (base<sup>-1</sup>) | Deletion rate (base<sup>-1</sup>) | Insertion rate (base<sup>-1</sup>) | Bases counted |
| --- | --- | --- | --- | --- | --- | --- |
| Mesophile | KleExo- | 10 | $4.20 \times 10^{-5}$ | $9.50 \times 10^{-7}$ | ND | 6,370,293 |
| | KleExo- | 20 | $6.91 \times 10^{-5}$ | $7.45 \times 10^{-7}$ | $5.31 \times 10^{-7}$ | 2,000,239 |
| | KleExo- | 37 | $1.07 \times 10^{-3}$ | ND | ND | 3,472,537 |
| | KleExo- | 50 | $1.38 \times 10^{-3}$ | $2.27 \times 10^{-6}$ | $2.01 \times 10^{-6}$ | 3,408,561 |
| | KleLF | 10 | $8.53 \times 10^{-6}$ | $2.18 \times 10^{-5}$ | $3.53 \times 10^{-6}$ | 5,822,486 |
| | KleLF | 20 | $1.56 \times 10^{-5}$ | ND | $8.32 \times 10^{-7}$ | 5,522,080 |
| | KleLF | 37 | $2.99 \times 10^{-5}$ | ND | ND | 4,317,523 |
| | KleLF | 50 | $2.00 \times 10^{-5}$ | ND | ND | 5,415,816 |
| | T4 | 10 | $3.25 \times 10^{-5}$ | $2.32 \times 10^{-6}$ | $4.81 \times 10^{-7}$ | 6,168,596 |
| | T4 | 37 | $5.92 \times 10^{-5}$ | $2.46 \times 10^{-6}$ | $4.84 \times 10^{-7}$ | 6,093,149 |
| | T4 | 50 | $1.45 \times 10^{-5}$ | ND | $5.11 \times 10^{-7}$ | 6,117,463 |

<sup>ND</sup>=error frequency below the detectable error baseline.

Intriguingly, we observe a substantial temperature dependence for psychrophilic and mesophilic polymerase substitution error rates. The effect is most dramatic on PIPI and KleExo-whose substitution rate rose by more than 5 and 30 fold, respectively, between reaction temperatures of 10°C and 50°C (**Tables 2–3**) (left and center panels in **Figures 4a**). Similarly, the deletion rate also scales positively with reaction temperature for PIPI, and KleExo- (**Tables 2–3**) (**Supplementary Figure 2b**). Curiously, the temperature effect on substitution error rate of KleLF and T4 polymerases is not monotonic, revealing a maximum substitution rate at 37°C (**Table 3**) (center panel in **Figures 4a**). This is consistent with the observation that T4 DNA polymerase’s error rate can decrease at a higher temperature ^39^. Comparison of KleLF and KleExo- suggests that the presence of exonuclease domain may have a dominant and inhibitory effect on error rate at high reaction temperature.

To investigate whether there is an activity-fidelity trade-off, we quantified the relationship between extension activity and substitution error rate for polymerases with and without exonuclease activity using linear regression model (**Figure 4b – Figure 4c**). For PIPI and KleLF, which are predicted to exhibit exonuclease activity, we found that extension activity (log_2_) significantly predicted substitution error rate (log_2_), *b* = 1.1502, *t*(14) = 4.385, *p* < .001. Strikingly, we observed a similar relationship in Taq and KleExo- polymerases which lack exonuclease activity, *b* = 1.1394, *t*(10) = 5.088, *p* < 0.01. Intuitively, our model predicts that for every doubling of polymerase activity, polymerase error rate increases by about a factor of 2.2. Extension activity (log_2_) also explained a significant portion of variance in substitution error rate (log_2_), *R*^2^ = 0.597, *F*(1, 13) = 19.23, *p* < .001 (**Figure 4b**), and *R*^2^ = 0.742, *F*(1, 9) = 25.89, *p* < .001 (**Figure 4c**). We next analyzed the changes in substitution error spectrum across reaction temperature. Reaction temperature had the most substantial effect on the substitution spectrum of KleExo-. As expected, an increase in reaction temperature is associated with an increased frequency of transversion relative to transition errors (**Supplemental Figure 2c**). Meanwhile, reaction temperature had a relatively mild effect on the substitution spectrum of other polymerases that we investigated.

### Distinct mutational profiles are driven by polymerase family type

Polymerase replication error rate is context-dependent ^40,41^. We wondered whether phylogenetically related polymerases that experienced vastly different temperature adaptation would produce substitution profiles similar to each other. We also asked whether these profiles varied across reaction temperature and polymerase phylogeny. To validate the phylogenetic relations, we aligned the primary peptide sequences of PIPI to polymerases with known family assignment (**Supplemental Figure 3a**). To analyze the relations between polymerase family and sequence context of substitution errors, we generated a 3-mer substitution spectrum for each condition. The entire spectrum consists of 192 permutations (4 possible nucleotides each at −1, 0, +1 positions, and 3 possible mutated bases). To enrich for informative motif, we kept 3-mer substitutions that occurred at greater than 20% frequency in at least a single reaction, and removed the rest from further analysis. This filtering step allows us to narrow down to 60 distinct 3-mer substitutions (∼31.3% of the entire 3-mer spectrum) and focus on substitutions that are common in at least some of the reactions. The hierarchical clustering of the filtered 3-mer substitution matrix reveals that substitution profiles generated by the same polymerase cluster across different reaction temperature. This indicates that the effect of reaction temperature on polymerase substitution error is independent of the sequence context (**Figure 5a**). Interestingly, we discover that the substitution profiles tended to cluster by phylogenetic distance of the polymerases in which substitutions generated by families A and B largely separated into distinct clusters regardless of their respective temperature adaptation range.

**Figure 5.**
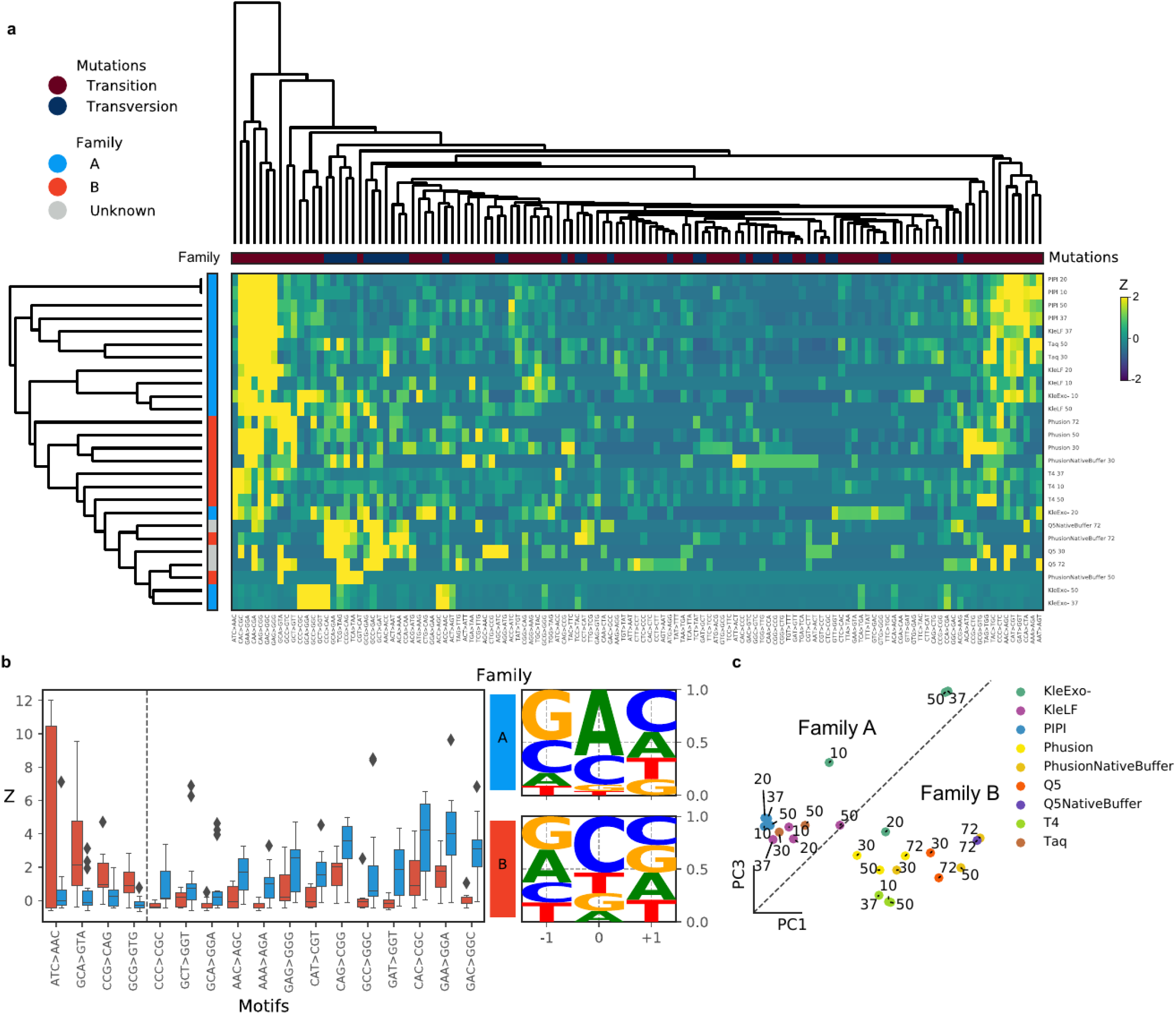
**(a)** Hierarchical clustering of the standardized 3-mer substitution rates naturally reveal clustering of error motif by the phylogenetic origins of polymerase. **(c)** Projection of polymerase reactions by principal components (PCs) 1 and 3 of standardized 3-mer substitution rates. **(b)** Substitution motifs that most distinguish errors made by family A and family B polymerases. Logo plots quantify the frequency of substitution motifs at −1, 0, and +1 positions that are generated by polymerase families A (top) or B (bottom).

To identify mutational profiles that best explain the observed substitution profiles, we trained a random forest decision tree to classify the polymerase family origin of each profile. By ranking the learned weight for each feature, we identified the most distinguishing mutational footprints for each polymerase family (**Figure 5b**). We discover that polymerase family A more frequently produced A → G transition when preceded by G/C at the −1 position. On the other hand, polymerase family B more frequently produced pyrimidine (C/T) → A transversion when preceded by a purine (A/G) at the −1 position. We then performed principal component analysis on the standardized the 3-mer error rate. This led us to identify principal components 1 and 3 that best explained the polymerase families’ substitution profiles. Overall, a high principal component 1 score is associated with the substitution profile of family B, while high principal component 3 score is associated with that of family A. Projection of the substitution profiles onto principal components 1 and 3 reveals a natural decision boundary on this space that separates substitution error spectrum between family A and B (**Figure 5c**). This suggests that conserved structural differences between the two polymerase families may underlie polymerase’s propensity to generate distinct mutational signatures independent of temperature adaptation experience and the reaction temperature.

## Discussion

DNA polymerases are the biological machinery underlying the transmission of genetic information and hold the key to many biotechnological innovations. While most studies have focused on polymerases of mesophilic or thermophilic origins, this is one of the first to provide a comprehensive biochemical characterization of activity and fidelity of bacterial DNA polymerases across a wide temperature adaptation range. We characterized the temperature-dependence of apparent replication activity in DNA polymerases. Earlier studies had reported that mesophilic and thermophilic DNA polymerases become increasingly inactivated at low temperature range ^34,36^. Consistent with their reports, we were unable to detect DNA replication activity of Klenow fragment at negative 19°C. Intriguingly, we detected robust activity with the psychrophilic DNA polymerase PIPI. To our knowledge, this is the first demonstration that DNA extension can be accomplished with a psychrophilic DNA polymerase below water’s freezing temperature (0°C). As these two polymerases are phylogenetic cousins, comparative study into the molecular mechanism for why the psychrophilic polymerase retains activity at low temperature may yield additional insights into the design principles of enzymes. Low temperature reaction is desirable as it can inhibit nuclease activity and reduce phototoxicity and DNA damages ^24,42^. Reactions in exotic environments such as the arctic and outer space may also require polymerases that can function at the low temperature range. The ability to engineer psychrophilic polymerases can thus enable nucleic acid amplification techniques directly on cryopreserved samples or in cold environment, reducing the risk of contamination and signal degradation ^43^. Combining polymerases adapted to different temperature range can, in principle, enable thermally multiplexed nucleic acid reaction that can substantially reduce assay time. One can also imagine combining the polymerases in a multi-step reaction assay, using temperature as a parameter to control reaction sequence. As secondary structures in nucleic acids are highly sensitive to thermal destabilization, psychrophilic polymerases may be used for detection of such structures. Recent reports also suggest that dynamic modulation of liquid-frozen phase can also provide a new niche for psychrophilic polymerase applications ^44,45^. The development of cold-adapted polymerases can greatly improve the capability of existing technologies and expand the temperature range of assays that one can perform.

In this study, we have also explored one of the long-standing questions in enzymology – is there a necessary trade-off between substrate specificity and activity? To address this question, we biochemically characterized bacterial DNA polymerase adapted to a wide range of environmental temperatures. Our results suggest that distinct temperature adaptation can differentially alter how the activity and fidelity of DNA polymerases change in response to environmental temperature fluctuations. We find that activity and fidelity have a log-linear relationship for many of the polymerases that we investigated. While the effect of temperature on Taq and T4 DNA polymerase error rate had been investigated ^39,46^, we present the most comprehensive report of temperature effect across polymerases on the single-molecule level, which enabled analysis of the temperature-dependent changes in the mutational spectrum. Our data shows that psychrophilic and mesophilic polymerases tend to be more heat-labile while thermophilic polymerases, with the exception of Taq, are largely heat resistant, consistent with previous reports ^47^. Furthermore, increased reaction temperature substantially increased the error rate of PIPI and *E. coli* DNA polymerase I (Klenow fragments) but not that of thermophilic polymerases such as Phusion and Q5. Nucleic acid analogues are of interest to a wide range of synthetic biology and biological engineering research as a means to expand orthogonal signals or building parts ^48^. This is often achieved through a combination of rational engineering and directed evolution on mesophilic or thermophilic polymerase templates ^49–52^. It is tempting to speculate that using cold-adapted polymerases as a starting template may achieve higher substrate promiscuity and thus greater incorporation rate of unnatural nucleotides; however, further experiments are required to validate this conjecture. Engineering of psychrophilic enzymes may help to improve most sequencing-by-synthesis sequencers available on the market as well as accelerate the application of nucleic acids analogues.

## Experimental Procedures

### Polymerase Cloning

*Psychromonas ingrahamii* 37 DNA Polymerases I (PIPI; IMG Gene ID: 639798289) sequence is provided by Joint Genome Institute, Integrated Microbial Genomes & Microbiomes (IMG/G) portal (IMG Taxon ID: 639633052). The PIPI sequence was codon-optimized for *E. coli* expression, synthesized as a gene fragment (1845 bp), and then cloned into pD454 T7 expression vector (<u>Ampicillin</u> resistance) by ATUM (formerly DNA 2.0) to generate a tandem fusion construct with maltose-binding protein (MBP) separated by a single HRV 3C cleavage site. Cloning product was first subcloned in NEB-5α strains and then purified using ZymoPURE plasmid miniprep kits (Zymo Research). The sequence of the expression construct was verified with Sanger sequencing by Sequetech. Plasmid map is also provided in **Supplemental file 1**.

### Polymerase Purification

Several expression and purification conditions were attempted to optimize the yield and purity of PIPI. In the end, we generated MBP-PIPI (112.4 kDa) fusion gene on a pMAL-c5x vector and expressed it in Arctic Express DE3 (Agilent Technologies), a bacterial expression cell line that expresses psychrophilic chaperones which promote proper protein folding during low temperature growth. Instead of LB, we grew the bacteria in a defined media M9ZB (**Supplemental file 2**) at 37°C until OD600 of 2.5 and induced expression with 1M IPTG at 10°C or 16°C for 20-24 hours. Cells were harvested in lysis buffer (20 mM Tris-HCl pH 7.50, 10% glycerol, 50 mM NaCl, 10 mM 2-Mercaptoethanol) with the additions of 400 U of DNase I and 1X protease inhibitor cocktail (No EDTA; ThermoFisher). Cells were lysed in French press homogenizer by passing through at 500 psi once, and at 10000 psi for three times. Clarified cell lysate was mixed with amylose resin at 4°C for three hours. The resin was then washed with 20 column volumes of chaperone removal buffer (20 mM Tris-HCl pH 7.50, 10% glycerol, 50 mM NaCl, 10 mM 2-Mercaptoethanol, 50 mM KCl, 5 mM ATP) and washed with 10 column volumes of lysis buffer (with no addition of DNase I or protease inhibitor). Bound proteins were eluted by incubating at 4°C for 3 hours with 10 mM of maltose. MBP-PIPI was subjected to HRV-3C cleavage at 4°C overnight and was further purified in HiLoad 16/60 Superdex 200 preparatory grade size exclusion column (GE Healthcare) to separate fractions of polymerase proteins from cleaved MBP and protease. Purified proteins were eluted in lysis buffer, concentrated in 30 kDa Amicon Ultra protein filters to ∼0.1 mg/mL, and then stored in −80°C until experiments.

### Protein quantification

Protein concentration was quantified by loading proteins on denaturing SDS-PAGE gel and stained with SYPRO Red following the manufacturer’s recommended protocol (ThermoFisher). Linear dynamic range of the quantification was established by separately staining a titration series of known amount of MBP-paramyosin-Sal. A secondary quantification measurement with Bioanalyzer Protein Analysis Kits (Agilent) provided similar concentration values as SYPRO Red assay (data not shown).

### Polymerase Activity Assay

Polymerase activity from 5°C to 50°C was measured with EvaEZ™ Fluorometric Polymerase Activity Assay Kit (Biotium) in the provided Tris buffer system based on the manufacturer’s protocol. Polymerase was diluted to a concentration that is saturated by substrate concentration. Briefly, assay mix and polymerase were equilibrated to the reaction temperature separately for 5 minutes before mixing. Fluorescence signals of Eva double-straned DNA (dsDNA) binding dye and ROX reference dye were measured in Bio-rad 96 well qPCR machine for 1 hour at constant reaction temperature with 30 seconds interval. A positive control sample with saturating amount of Klenow Exo- polymerase was included for each experiment to ensure that maximal fluorescence signals remained constant throughout measurement. The initial slope of the fluorescence gain for each polymerase was used to estimate the replication’s steady-state speed.

### Gel Extension Assay

For measurement of subzero polymerase activity, we prepared 0.5 μL of polymerase pre-diluted to 5 nM in MOPS buffer (pH 8.50). In parallel, we also prepared 9.5 μL of primed extension mix containing a final concentration in 10 μL volume of 10 nM primed template (TGATG**GCGCCGTGACAGTGAAT**) with 5’-ATTOTM 633 fluorescent label (/5ATTO633N/**ATTCACTGTCACGGCGC**; iDT) (**Supplemental file 1**), 50 μM dNTP, 10 mM MOPS pH 8.50, 30% glycerol, 1.5 mM MgCl2, 0.1 mg/mL BSA, 50 mM KCl. Reaction master mix and polymerase were separately equilibrated to −19°C on a TropiCoolerTM (Boekel Scientific) for 30 minutes, and then rapidly mixed using pre-chilled pipette tips. One microliter aliquot of the reaction was quenched in 9 μL of 90% formamide with 50 mM EDTA pH 8.0 and heat-denatured at 95°C for 5 minutes before cooling on ice. Gels were imaged on Typhoon 9410 imager (GE).

### Molecular Barcoding and Library Preparation

We purified pUC19 plasmid template from a single, transformed clone of NEB-5α (New England Biolabs) grown in LB and 100 μg/mL ampicillin antibiotic. The plasmid’s spontaneous mutagenesis rate is around ∼10^−7^ – 10^−8^ ^27,28^, which is below the error rate of Q5 polymerase that we used for second-strand synthesis. About one μg of purified pUC19 was co-digested with ten units of SapI and NdeI at 37°C for 24 minutes to yield 509 bp fragment. Digested fragment was purified twice with 0.35X (V:V) of Ampure XP beads to remove backbone vector and restriction enzymes. Concentration and purity of the co-digested pUC19 template were determined on Bioanalyzer using a high-sensitivity dsDNA quantification kit. Sequencing error in Illumina short-read sequencing occurs at a much higher frequency (∼10^−2^ to 10^−3^ per base) than polymerase replication error (10^−3^ to 10^−7^ per base). In order to confidently call out replication errors, sequencing errors have to be distinguished from replication errors.

We incorporated Unique Molecular Indices (UMIs), a 15 bp randomer, into each primer that serves as a molecular barcode ^29^ (**Supplemental file 1**). This barcode is used to resolve the molecular origin of each sequenced read. As each molecule is sequenced multiple times, consensus sequences can be constructed to infer true replication errors. All reactions were conducted in a buffer made up of 10 mM MOPS pH 8.50, 30% glycerol, 1.5 mM MgCl2, 0.1 mg/mL BSA, 50 mM KCl unless otherwise stated. We used MOPS buffer instead of Tris-based buffer as it has a much smaller ΔpKa/ΔT scaling which minimizes changes in buffer pH at different temperatures. UMI index primer and pUC19 template were mixed at 100:1 molar ratio, denatured at 95°C for 30 seconds, and annealed at 52°C for 2 minutes. We then pre-incubated the reaction mixture in 96 well plate at the desired reaction temperature for 2 minutes before the addition of polymerase (volume of reaction mixture to volume of polymerase is 19:1). Reaction time for each polymerase and condition was determined using fluorometric activity assays and qPCR measurement. The reaction was quenched with 50 mM EDTA and then purified with 1X Ampure XP beads. We then synthesized the complementary strand using the reverse UMI primer and Q5 DNA polymerase, a high-fidelity polymerase, in its native reaction buffer (NEB) by incubating at 72°C for 10 minutes, which is followed by EDTA quenching. We performed a second round of DNA purification with 1X Ampure XP beads and quantified molecular concentration of the resulting products using qPCR. A minimum of 20000 barcoded molecules from each reaction was amplified in two rounds of 13 PCR cycles, with Ampure XP purification between each successive round of amplification ^29^. Agilent Bioanalyzer dsDNA kit was used to assess the purity of the amplified product. Some reactions were repeated multiple times in order to obtain sufficient coverage for the analysis.

### Replication Error Analysis

The library was sequenced on MiSeq using V3 2 × 300 bp chemistry kit to yield approximately 25 million reads. We sequenced some libraries multiple times to ensure sufficient sequencing coverage for the analysis. Each read is sorted into groups according to reaction indices. Each read is trimmed to a minimum of 150 length and Q score of 20 (Phred+33 quality score), ensuring that the composite reads will cover the entire template sequence. Reads with the same reaction indices are then grouped according to their UMI barcodes. Reads originating from the same replicated molecule would share the same UMI barcode. The consensus for each position was determined by 90% majority within the UMI family that shares the same base. Bases with less than 90% majority consensus may contain PCR-induced errors and thus are not included for downstream analysis. Consensus sequences with non-matching bases in the first five positions are removed to avoid analysis of mis-primed products. We observed that the apparent error rate for a polymerase decreased and plateaued as the consensus number increased, and we picked a minimum of 5 or 10 consensus threshold for each molecule to improve accuracy in replication error calling. The consensus sequence from each UMI family is then aligned to the reference sequence using BWA-MEM ^30^. Substitution, insertion, and deletion errors are determined by comparing each UMI molecule’s consensus sequence to the pUC19 reference sequence without counting ambiguous assignments. We did not observe any pre-existing mutation in the pUC19 reference that occurred across all reactions.

## Data Availability

Processed data are available as Supplemental files (Supplemental files 3-7). Raw and intermediate fastq files and supplemental files are deposited on Dryad: Xue, Yuan; Braslavsky, Ido; Quake, Stephen (2021), Temperature Effect on Polymerase Fidelity, Dryad, Dataset, https://doi.org/10.5061/dryad.76hdr7stv. Link to download the data: https://datadryad.org/stash/share/vE81mjJcahAkgiIed7prnOmMNe9NIa4mgiDCDcz4F3w.

## Acknowledgments

YX thanks professors James Berger and Daniel Herschlag for helpful advice and discussions. YX thanks Richard Pfuetzner, Steven Wilson, Qiangjun Zhou, and Austin Wang for sharing equipment and suggestions. YX thanks Norma Neff, Fabio Zanini, Felix Horns, Derek Croote, Mark Kowarsky, Brian Yu, Bojk Berghuis, and Nimit Jain for discussions and suggestions. YX is supported by the Stanford Interdisciplinary Graduate Fellowship and Weiland Family Fellowship. IB acknowledges support from Stanford University and from The Hebrew University of Jerusalem. SRQ is supported by the Chan Zuckerberg Biohub. This project is also supported by the Templeton Foundation.

---

**Supplemental figure 1.**
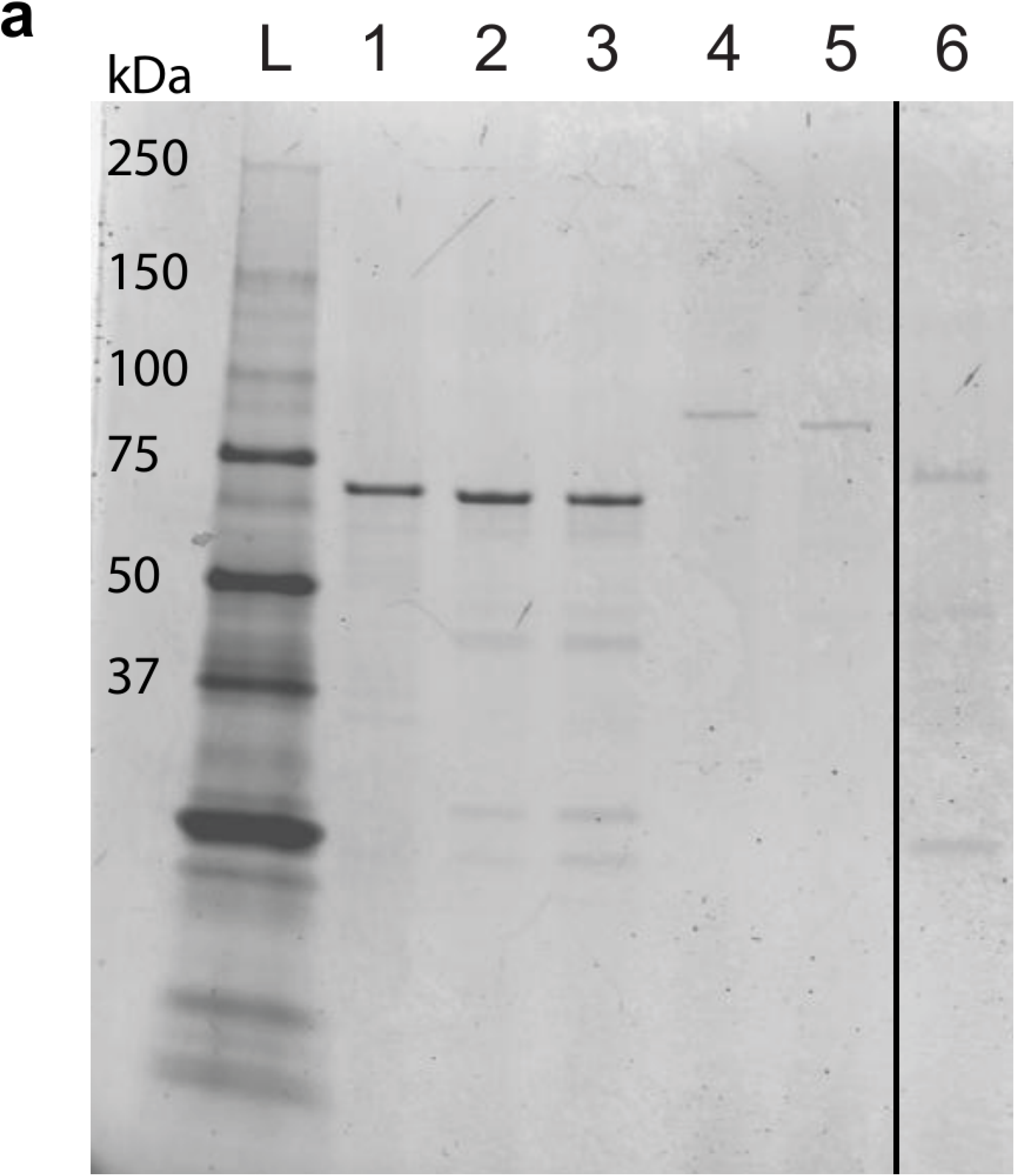
**(a)** SDS-PAGE analysis of purified DNA polymerases. Lane L: PrecisionPlus Ladder (Bio-rad). Lane 1: MBP-paramyosin protein control (NEB). Lane 2: Klenow Exo-(NEB). Lane 3: Klenow LF (NEB). Lane 4: Q5 (NEB). Lane 5: Taq (NEB). Lane 6: PIPI (69.9 kDa).

**Supplemental figure 2.**
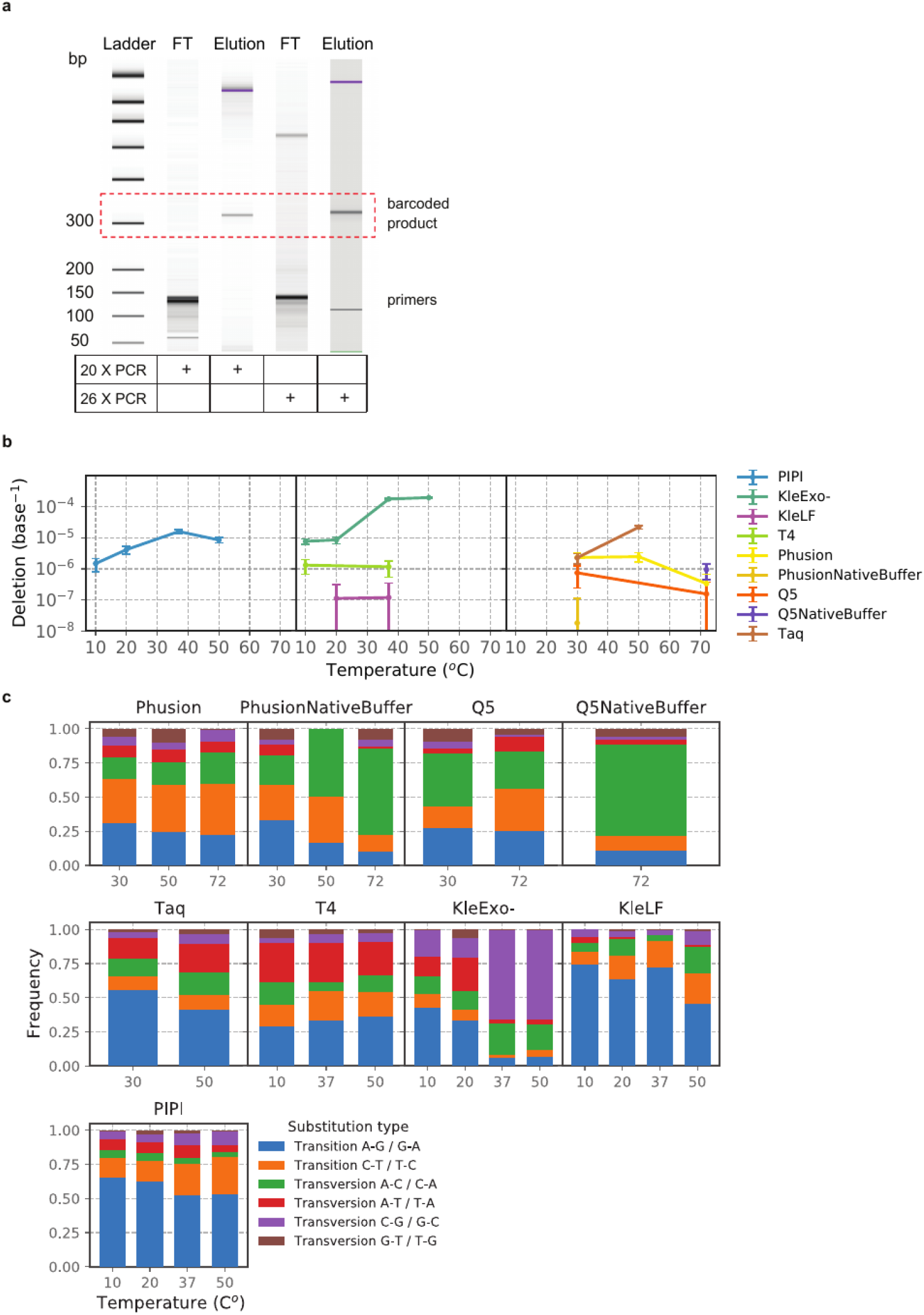
**(a)** High-sensitivity Bioanalyzer DNA (Agilent) assay of replicated products after 20 or 26 rounds of PCR amplification in the flow-through (FT) or elution fractions of Ampure purification output. **(b)** Deletion error rates (per base) of psychrophilic (left), mesophilic (center), and thermophilic (right) DNA polymerases as a function of reaction temperature. Data points with rates below the Q5 polymerase deletion rate baseline are excluded. **(c)** Substitution spectrum for all DNA polymerases tested in this study. The height of each colored bar reflects the frequency of a particular substitution type in all the substitution events observed for a polymerase reaction at the specified reaction temperature.

**Supplemental figure 3.**
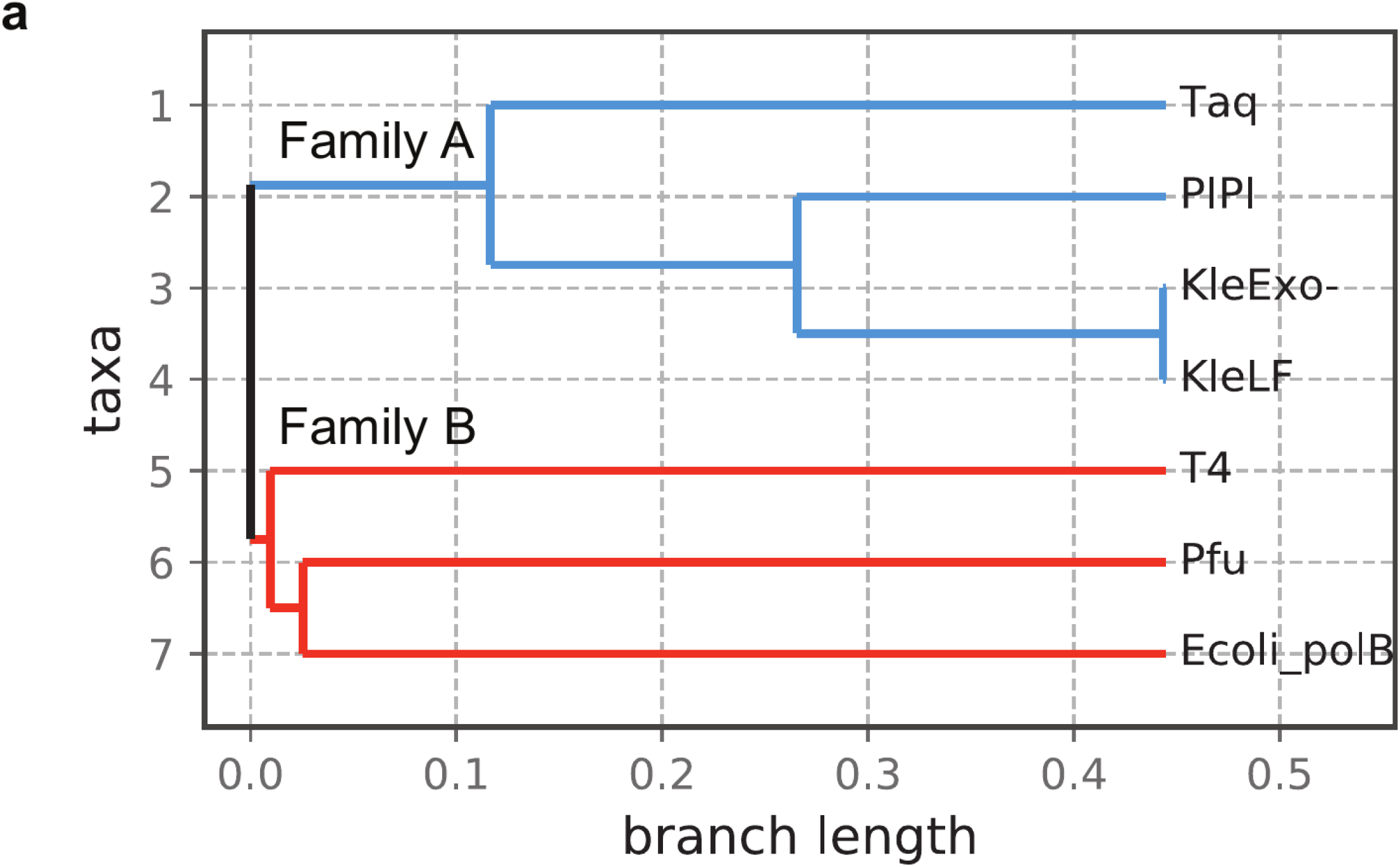
**(a)** Multiple sequence alignment (Clustal Omega) of peptide sequences for polymerases used in this study. While Phusion DNA polymerase sequence is undisclosed, it is expected to be a derivative of Pyrococcus DNA polymerase II (Pfu). Family A cluster consists of Taq, PIPI, and Klenow polymerases. Family B polymerases consist of T4, Pfu, and *E. coli* DNA polymerase II (Ecoli_polB).

## References

1. Ratkowsky, D. A., Olley, J., McMeekin, T. A. & Ball, A. Relationship between temperature and growth rate of bacterial cultures. J. Bacteriol. 149, 1–5 (1982).

2. Ingraham, J. L. & Stokes, J. L. Distinguished By. Bacteriolody Rev. 23, 97–108 (1934).

3. Svingor, Á., Kardos, J., Hajdú, I., Németh, A. & Závodszky, P. A better enzyme to cope with cold: Comparative flexibility studies on psychrotrophic, mesophilic, and thermophilic IPMDHS. J. Biol. Chem. 276, 28121–28125 (2001).

4. Ingraham, J. & Bailey, G. Comparative study of effect of temperature on metabolism of psychrophilic and mesophilic bacteria. J. Bacteriol. 77, 609–613 (1959).

5. Wintrode, P. L., Miyazaki, K. & Arnold, F. H. Cold adaptation of a mesophilic subtilisin-like protease by laboratory evolution. J. Biol. Chem. 275, 31635–31640 (2000).

6. Lonhienne, T., Gerday, C. & Feller, G. Psychrophilic enzymes: Revisiting the thermodynamic parameters of activation may explain local flexibility. Biochim. Biophys. Acta - Protein Struct. Mol. Enzymol. 1543, 1–10 (2000).

7. Lonhienne, T. et al. Modular structure, local flexibility and cold-activity of a novel chitobiase from a psychrophilic antarctic bacterium. J. Mol. Biol. 310, 291–297 (2001).

8. Liang, Z. X. et al. Evidence for increased local flexibility in psychrophilic alcohol dehydrogenase relative to its thermophilic homologue. Biochemistry 43, 14676–14683 (2004).

9. Seifert, M. et al. Temperature controlled high-throughput magnetic tweezers show striking difference in activation energies of replicating viral RNA-dependent RNA polymerases. Nucleic Acids Res. 48, 5591–5602 (2020).

10. Miyazaki, K., Wintrode, P. L., Grayling, R. A., Rubingh, D. N. & Arnold, F. H. Directed evolution study of temperature adaptation in a psychrophilic enzyme. J. Mol. Biol. 297, 1015–1026 (2000).

11. Nguyen, V. et al. Evolutionary drivers of thermoadaptation in enzyme catalysis. Science (80-. ). 355, 289–294 (2017).

12. Tsigos, I., Velonia, K., Smonou, I. & Bouriotis, V. Purification and characterization of an alcohol dehydrogenase from the Antarctic psychrophile Moraxella sp. TAE123. Eur. J. Biochem. 254, 356–362 (1998).

13. D’Amico, S., Sohier, J. S. & Feller, G. Kinetics and Energetics of Ligand Binding Determined by Microcalorimetry: Insights into Active Site Mobility in a Psychrophilic α-Amylase. J. Mol. Biol. 358, 1296–1304 (2006).

14. Loeb, L. A. & Monnat, R. J. DNA polymerases and human disease. Nat. Rev. Genet. 9, 594–604 (2008).

15. Sierra, H., Cordova, M., Chen, C. S. J. & Rajadhyaksha, M. Confocal imaging-guided laser ablation of basal cell carcinomas: An ex vivo study. J. Invest. Dermatol. 135, 612–615 (2015).

16. Broughton, B. C. et al. Molecular analysis of mutations in DNA polymerase η in xeroderma pigmentosum-variant patients. Proc. Natl. Acad. Sci. U. S. A. 99, 815–820 (2002).

17. Donigan, K. A. et al. Human POLB gene is mutated in high percentage of colorectal tumors. J. Biol. Chem. 287, 23830–23839 (2012).

18. Cabelof, D. C. et al. Haploinsufficiency in DNA polymerase β increases cancer risk with age and alters mortality rate. Cancer Res. 66, 7460–7465 (2006).

19. Aschenbrenner, J. & Marx, A. DNA polymerases and biotechnological applications. Curr. Opin. Biotechnol. 48, 187–195 (2017).

20. Dean, F. B., Nelson, J. R., Giesler, T. L. & Lasken, R. S. Rapid amplification of plasmid and phage DNA using Phi29 DNA polymerase and multiply-primed rolling circle amplification. Genome Res. 11, 1095–1099 (2001).

21. Chen, M. et al. Comparison of Multiple Displacement Amplification (MDA) and Multiple Annealing and Looping-Based Amplification Cycles (MALBAC) in single-cell sequencing. PLoS One 9, 1–12 (2014).

22. Schmitt, M. W. et al. Detection of ultra-rare mutations by next-generation sequencing. Proc. Natl. Acad. Sci. U. S. A. 109, 14508–14513 (2012).

23. Smith, T., Heger, A. & Sudbery, I. UMI-tools: Modeling sequencing errors in Unique Molecular Identifiers to improve quantification accuracy. Genome Res. 27, 491–499 (2017).

24. Potapov, V. & Ong, J. L. Examining sources of error in PCR by single-molecule sequencing. PLoS One 12, 1–19 (2017).

25. Schleper, C., Swanson, R. V., Mathur, E. J. & Delong, E. F. Characterization of a DNA polymerase from the uncultivated psychrophilic archaeon Cenarchaeum symbiosum. J. Bacteriol. 179, 7803–7811 (1997).

26. Piotrowski, Y., Gurung, M. K. & Larsen, A. N. Characterization and engineering of a DNA polymerase reveals a single amino-acid substitution in the fingers subdomain to increase strand-displacement activity of A-family prokaryotic DNA polymerases. BMC Mol. Cell Biol. 20, 1–11 (2019).

27. Deatherage, D. E., Leon, D., Rodriguez, Á. E., Omar, S. K. & Barrick, J. E. Directed evolution of Escherichia coli with lower-than-natural plasmid mutation rates. Nucleic Acids Res. 46, 9236–9250 (2018).

28. Camps, M., Naukkarinen, J., Johnson, B. P. & Loeb, L. A. Targeted gene evolution in Escherichia coli using a highly error-prone DNA polymerase I. Proc. Natl. Acad. Sci. U. S. A. 100, 9727–9732 (2003).

29. Lee, D. F., Lu, J., Chang, S., Loparo, J. J. & Xie, X. S. Mapping DNA polymerase errors by single-molecule sequencing. Nucleic Acids Res. 44, e118 (2016).

30. Li, H. Aligning sequence reads, clone sequences and assembly contigs with BWA-MEM. 00, 1–3 (2013).

31. Breezee, J., Cady, N. & Staley, J. T. Subfreezing growth of the sea ice bacterium ‘Psychromonas ingrahamii’. Microb. Ecol. 47, 300–304 (2004).

32. Auman, A. J., Breeze, J. L., Gosink, J. J., Kämpfer, P. & Staley, J. T. Psychromonas ingrahamii sp. nov., a novel gas vacuolate, psychrophilic bacterium isolated from Arctic polar sea ice. Int. J. Syst. Evol. Microbiol. 56, 1001–1007 (2006).

33. Riley, M. et al. Genomics of an extreme psychrophile, Psychromonas ingrahamii. BMC Genomics 9, 1–19 (2008).

34. Brown, H. S. & Licata, V. J. Enthalpic switch-points and temperature dependencies of DNA binding and nucleotide incorporation by Pol i DNA polymerases. Biochim. Biophys. Acta - Proteins Proteomics 1834, 2133–2138 (2013).

35. Datta, K., Wowor, A. J., Richard, A. J. & LiCata, V. J. Temperature dependence and thermodynamics of klenow polymerase binding to primed-template DNA. Biophys. J. 90, 1739–1751 (2006).

36. Rentergent, J., Driscoll, M. D. & Hay, S. Time Course Analysis of Enzyme-Catalyzed DNA Polymerization. Biochemistry 55, 5622–5634 (2016).

37. Ling, L. L., Keohavong, P., Dias, C. & Thilly, W. G. Optimization of the polymerase chain reaction with regard to fidelity: Modified T7, Taq, and vent DNA polymerases. Genome Res. 1, 63–69 (1991).

38. Altschuler, S. M. What is LBS and What Does It Mean „ for Your Business? Doctor vol. 3 26–28 (2010).

39. Bessman, M. J. & Reha-Krantz, L. J. Studies on the Biochemical Basis of Spontaneous Mutation - Effect of Temperature on Mutation Frequency. J. Mol. Biol. 115–123 (1977).

40. de Paz, A. M. et al. High-resolution mapping of DNA polymerase fidelity using nucleotide imbalances and next-generation sequencing. Nucleic Acids Res. 46, e78 (2018).

41. Schroeder, J. W., Hirst, W. G., Szewczyk, G. A. & Simmons, L. A. The Effect of Local Sequence Context on Mutational Bias of Genes Encoded on the Leading and Lagging Strands. Curr. Biol. 26, 692–697 (2016).

42. Adam, M., Potter, A. S. & Potter, S. S. Psychrophilic proteases dramatically reduce single-cell RNA-seq artifacts: A molecular atlas of kidney development. Development 144, 3625–3632 (2017).

43. Guillaumet-Adkins, A. et al. Single-cell transcriptome conservation in cryopreserved cells and tissues. Genome Biol. 18, 1–15 (2017).

44. Glen, J. W. Ice Physics. Physics Bulletin vol. 37 (Oxford University Press, 1986).

45. Preis, S. G. et al. Labyrinth ice pattern formation induced by near-infrared irradiation. Sci. Adv. 5, (2019).

46. Tindall, K. R. & Kunkel, T. A. Fidelity of DNA Synthesis by the Thermus aquaticus DNA Polymerase. Biochemistry 27, 6008–6013 (1988).

47. Siddiqui, K. S. & Cavicchioli, R. Cold-adapted enzymes. Annu. Rev. Biochem. 75, 403–433 (2006).

48. Hirao, I. et al. An unnatural hydrophobic base pair system: Site-specific incorporation of nucleotide analogs into DNA and RNA. Nat. Methods 3, 729–735 (2006).

49. Gardner, A. F. et al. Therminator DNA Polymerase: Modified nucleotides and unnatural substrates. Front. Mol. Biosci. 6, 1–9 (2019).

50. Chen, T. & Romesberg, F. E. Directed polymerase evolution. FEBS Lett. 588, 219–229 (2014).

51. Xia, G. et al. Directed evolution of novel polymerase activities: Mutation of a DNA polymerase into an efficient RNA polymerase. Proc. Natl. Acad. Sci. U. S. A. 99, 6597–6602 (2002).

52. Laos, R., Thomson, J. M. & Benner, S. A. DNA polymerases engineered by directed evolution to incorporate nonstandard nucleotides. Front. Microbiol. 5, 1–14 (2014).

