## Supplementary Files 1 - 8 for "Temperature Effect on Polymerase Fidelity": sfile_1_plasmid_primer_sequences.pdf

**Supplemental text 1**

pD454-PIPI (7096 bases, circular construct):

ctgggcggttctgataacgagtaatcgttaatccgcaaataacgtaaaaacccgcttcggcgggtttttttatgggggggagtttagggaaagagcatttgtcagaatatttaagggcgcc
tgtcactttgcttgatatatgagaattatttaaccttataaatgagaaaaaagcaacgcactttaaataagatacgttgctttttcgattgatgaacacctataattaaactattcatctattattta
tgatttttgtatatacaatatttctagtttgttaaagagaattaagaaaataaatctcgaaaataataaagggaaaatcagtttttgatatcaaaattatacatgtcaacgataatacaaaatat
aatacaaactataagatgttatcagtatttattatcatttagaataaattttgtgtcgcccttccgcgaaattaatacgactcactataggggaattgtgagcggataacaattcccctctaga
aataattttgtttaactttttgagaccttaaagaaggagatataaaATGCATCATCACCACCATCATAAAACTGAAGAAGGTAAACTGGTAAT
CTGGATTAACGGCGATAAAGGCTATAACGGTTTGGCTGAAGTCGGTAAGAAATTCGAGAAAGATACCGGAATTA
AAGTCACCGTTGAGCATCCGGATAAACTGGAAGAGAAATTCCCACAGGTTGCGGCAACTGGCGATGGCCCTGA
CATTATCTTCTGGGCACACGACCGCTTTGGTGGCTACGCTCAATCTGGCCTGTTGGCTGAAATCACCCCGGACA
AAGCGTTCCAGGACAAGCTGTATCCGTTTACCTGGGATGCCGTACGTTACAACGGCAAGCTGATTGCTTACCCG
ATCGCTGTTGAAGCGTTATCGCTGATTTATAACAAAGATCTGCTGCCGAACCCGCCAAAAACCTGGGAAGAGAT
CCCGGCGCTGGATAAAGAACTGAAAGCGAAAGGTAAGAGCGCGCTGATGTTCAACCTGCAAGAACCGTACTT
CACCTGGCCGCTGATTGCTGCTGACGGGGGTTATGCGTTCAAGTATGAAAACGGCAAGTACGACATTAAAGAC
GTGGGCGTGGATAACGCTGGCGCGAAAGCGGGTCTGACCTTCCTGGTTGACCTGATTAAAAACAAACACATGA
ATGCAGACACCGATTACTCCATCGCAGAAGCTGCCTTTAATAAAGGCGAAACAGCGATGACCATCAACGGCCC
GTGGGCATGGTCCAACATCGACACCAGCAAAGTGAATTATGGTGTAACGGTACTGCCGACCTTCAAGGGTCAA
CCATCCAAACCGTTCGTTGGCGTGCTGAGCGCAGGTATTAACGCCGCCAGTCCGAACAAAGAGCTGGCAAAAG
AGTTCCTCGAAAACTATCTGCTGACTGATGAAGGTCTGGAAGCGGTTAATAAAGACAAACCGCTGGGTGCCGT
AGCGCTGAAGTCTTACGAGGAAGAGTTGGCGAAAGATCCACGTATTGCCGCCACTATGGAAAACGCCCAGAA
AGGTGAAATCATGCCGAACATCCCGCAGATGTCCGCTTTCTGGTATGCCGTGCGTACTGCGGTGATCAACGCCG
CCAGCGGTCGTCAGACTGTCGATGAAGCCCTGAAAGACGCGCAGACTAATCTGGAAGTTCTTTTTAATTCGAG
CTCGAACAACAACAACAATAACAATAACAACAACCTCGGGATCGAGGGAAGGATTTCACATATGCTGGAAGTT
CTTTTTCAGGGTCCT**ATGATCGATCGTAGCGGCTACAAAACCATCTATACGCAAGGCCAACTGACCCAGTG**
**GATCGCGAAGTTAGAGGCCGCTGAGCTGTTTAGCTTCGATACCGAAACCACCTCGTTGGACTACATGCA**
**AGCGCGTATCGTGGGCTTGAGCTTTGCGGTTCAGAGCAATGTTGGTAGCGATGAGCAGCCGATTATCGA**
**AGCCGCCTATCTGCCACTGGCCCACGACTACATCGATGCACCGAAACAGCTCGATCTGACGACCACTCT**
**GGAGAAACTGCGCCCGCTGCTGGAGTCTGAGAAATATAAAAAAGTCGGTCAGAACCTGAAATATGACCG**
**TAGCGTCTTGCTGAATCACGGTATCGAGCTGAAGGGTATTAAGTTCGACACGATGCTGGAGAGCTATGT**
**GCTGGATTCCACCGGTCGTCATGACATGGACACCCTGGCGTTAAAGTACCTGGGCCATCAGTGCATCAG**
**CTTCGAAGAAATTGCTGGTAAGGGTAAAAAGCAACTGCCGTTCAATCAGATCTCCATCATTGAAGCGGC**
**ACCGTACGCGGCGGAAGATGCAGATGTTACGTTGCGTCTGCACTTGCAGTTGTTTGAGCAACTGAGCGC**
**TGAGCCGAAACTTCTGAGCGTGTTTAACAATATCGAGCTGCCGCTGTTGACGGTCCTGTCCGATATTGA**
**GCGTGGTGGCGTTGAGATTGATTCGGACCTGTTGACGAAACAAAGCGCGGAAATTGGCAAGCGTCTGC**
**TGGAACTGGAAGCACTGGCTTATGAAGAGGCCGGCAAACCGTTCAACCTGAGCTCTCCGAAACAGCTG**
**CAGACCATTCTGTACGATGAGCTGGAACTGCCGGTTCTGAAGAAAACCCCGAAGGGTGCGCCGAGCAC**
**CGCCGAGGAAGTGCTGCAAGAACTGGCATTGACGTACCCTCTGCCAAAGCTGATCATCGAGCACCGTG**
**GTCTG12AGCAAACTGAAAAGCACCTATACGGACAAGCTGCCGAAAATGGTTAATGAAAAGACTGGCCG**
**TCTGCACACCAGCTATCAACAAGCCGTGACGGTGACGGGTCGCCTGTCCTCTACGGACCCGAATCTGCA**
**GAATATCCCGATTCGCTCTGCAGAGGGCCGTCGTATCCGTCAAGCGTTCATTGCGCAGCCTGGTTACAA**
**GATCGTCGCGGCGGACTACAGCCAAATCGAGTTGCGTATCATGGCGCATCTGAGCCAGGACAAGGGCC**
**TGCTGGACGCATTCTCCACCGGCAAAGACATTCACAAAGCAACCGCGAGCGAAGTTTTCAGCGTTCCGC**
**TGGATGAAGTCACTACCGAGCAGCGCCGTTCCGCGAAGGCTATCAACTTTGGCTTGATCTATGGCATGA**
**GCGCGTTCGGCCTGGCGAAGCAACTCAACATTGGTCGCCACGAAGCACAGCTGTATATGGACAAATATT**
**TTAACCGTTACCCGGGTGTGCTGATTTACATGGAAGATACCCGCAGCCTGGCGAACGAAAAGGCTTACG**
**TTGAAACCATTCTGGGTAGACGTTTGCAGCTGCCGAATATTAAGAGCCGCAATGGTATGCTGAAAAAAG**
**CCGCGGAGCGTGCCGCGATTAACGCGCCGATGCAAGGTACCGCTGCGGATATTATCAAAAAAGCAATGA**
**TTGACATGGCAGACTGGATTGCGCAGAAGTCCCCGGGTAGCGTTCAAATGCTGATGCAAGTGCATGATG**
**AACTGGTCTTTAGCATTAAGGAAGAATTGGTCGAGAGCTACACGAAAGAAATTCAGGCCATCATGGCAA**
**AAGCCGCAGATCTGGATGTGCCACTGATCGCTGACGCCGGTGTCGGTGATAACTGGGACGAAGCGCAC**
**TAA**ggttagataatagggtctcacccccctagcataacccccttggggcctctaaacgggtcttgaggggttttttgcccctgagacgcgtcaatcgagttcgtacctaagggcgacac
cccctaattagcccgggcgaaaggcccagtctttcgactgagcctttcgttttatttgatgcctggcagttccctactctcgcatggggagtccccacactaccatcggcgctacggcgt
ttcacttctgagttcggcatggggtcaggtgggaccaccgcgctactgccgccaggcaaacaaggggtgttatgagccatattcaggtataaatgggctcgcgataatgttcagaatt
ggttaattggttgtaacactgacccctatttgtttatttttctaaatacattcaaatatgtatccgctcatgagacaataaccctgataaatgcttcaataatattgaaaaaggaagaatatgagt
attcaacatttccgtgtcgcccttattcccttttttgcggcattttgccttcctgtttttgctcacccagaaacgctggtgaaagtaaaagatgctgaagatcagttgggtgcacgagtgggtt
acatcgaactggatctcaacagcggtaagatccttgagagttttcgccccgaagaacgttttccaatgatgagcacttttaaagttctgctatgtggcgcggtattatcccgtattgacgc
cgggcaagagcaactcggtcgccgcatacactattctcagaatgacttggttgagtactcaccagtcacagaaaagcatcttacggatggcatgacagtaagagaattatgcagtgct

gccataaccatgagtgataacactgcggccaacttacttctgacaacgatcggaggaccgaaggagctaaccgcttttttgcacaacatgggggatcatgtaactcgccttgatcgttg
ggaaccggagctgaatgaagccataccaaacgacgagcgtgacaccacgatgcctgtagcgatggcaacaacgttgcgcaaactattaactggcgaactacttactctagcttccc
ggcaacaattaatagactggatggaggcggataaagttgcaggaccacttctgcgctcggcccttccggctggctggtttattgctgataaatccggagccggtgagcgtggttctcg
cggtatcatcgcagcgctggggccagatggtaagccctcccgtatcgtagttatctacacgacggggagtcaggcaactatggatgaacgaaatagacagatcgctgagataggtg
cctcactgattaagcattggtaagcggcgcgccatcgaatggcgcaaaacctttcgcggtatggcatgatagcgcccggaagagagtcaattcagggtggtgaatatgaaaccagt
aacgttatacgatgtcgcagagtatgccggtgtctcttatcagaccgtttcccgcgtggtgaaccaggccagccacgtttctgcgaaaacgcgggaaaaagtggaagcggcgatgg
cggagctgaattacattcccaaccgcgtggcacaacaactggcgggcaaacagtcgttgctgattggcgttgccacctccagtctggccctgcacgcgccgtcgcaaattgtcgcg
gcgattaaatctcgcgccgatcaactgggtgccagcgtggtggtgtcgatggtagaacgaagcggcgtcgaagcctgtaaagcggcggtgcacaatcttctcgcgcaacgcgtca
gtgggctgatcattaactatccgctggatgaccaggatgccattgctgtggaagctgcctgcactaatgttccggcgttatttcttgatgtctctgaccagacacccatcaacagtattatt
ttctcccatgaggacggtacgcgactgggcgtggagcatctggtcgcattgggtcaccagcaaatcgcgctgttagcgggcccattaagttctgtctcggcgcgtctgcgtctggctg
gctggcataaatatctcactcgcaatcaaattcagccgatagcggaacgggaaggcgactggagtgccatgtccggttttcaacaaaccatgcaaatgctgaatgagggcatcgttc
ccactgcgatgctggttgccaacgatcagatggcgctgggcgcaatgcgcgccattaccgagtccgggctgcgcgttggtgcggatatctcggtagtgggatacgacgataccga
agatagctcatgttatatcccgccgttaaccaccatcaaacaggattttcgcctgctggggcaaaccagcgtggaccgcttgctgcaactctctcagggccaggcggtgaagggcaa
tcagctgttgccagtctcactggtgaaaagaaaaaccaccctggcgcccaatacgcaaaccgcctctccccgcgcgttggccgattcattaatgcagctggcacgacaggtttcccg
actggaaagcgggcagtgactcatgaccaaaatcccttaacgtgagttacgcgcgcgtcgttccactgagcgtcagaccccgtagaaaagatcaaaggatcttcttgagatcctttttt
tctgcgcgtaatctgctgcttgcaaacaaaaaaaccaccgctaccagcggtggtttgtttgccggatcaagagctaccaactctttttccgaaggtaactggcttcagcagagcgcaga
taccaaatactgttcttctagtgtagccgtagttagcccaccacttcaagaactctgtagcaccgcctacatacctcgctctgctaatcctgttaccagtggctgctgccagtggcgataa
gtcgtgtcttaccgggttggactcaagacgatagttaccggataaggcgcagcggtcgggctgaacggggggttcgtgcacacagcccagcttggagcgaacgacctacaccga
actgagatacctacagcgtgagctatgagaaagcgccacgcttcccgaagggagaaaggcggacaggtatccggtaagcggcagggtcggaacaggagagcgcacgaggga
gcttccagggggaaacgcctggtatctttatagtcctgtcgggtttcgccacctctgacttgagcgtcgatttttgtgatgctcgtcaggggggcggagcctatggaaaaacgccagca
acgcggcctttttacggttcctggccttttgctggccttttgctcacatgttctttcctgcgttatcccctgattctgtggataaccgtattaccgcctttgagtgagctgataccgctcgccgc
agccgaacgaccgagcgcagcgagtcagtgagcgaggaagcggaaggcgagagtagggaactgccaggcatcaaactaagcagaaggcccctgacggatggcctttttgcgt
ttctacaaactctttctgtgttgtaaaacgacggccagtcttaagctcgggcccc

*upper cases: open reading frame of expression construct.
*upper cases + **boldened**: coding sequence for PIPI.

pUC19 (2686 bases, circular construct):

gacgaaagggcctcgtgatacgcctatttttataggttaatgtcatgataataatggtttcttagacgtcaggtggcacttttcggggaaatgtgcgcggaacccctatttgtttatttttcta
aatacattcaaatatgtatccgctcatgagacaataaccctgataaatgcttcaataatattgaaaaaggaagagtatgagtattcaacatttccgtgtcgcccttattcccttttttgcggca
ttttgccttcctgtttttgctcacccagaaacgctggtgaaagtaaaagatgctgaagatcagttgggtgcacgagtgggttacatcgaactggatctcaacagcggtaagatccttgag
agttttcgccccgaagaacgttttccaatgatgagcacttttaaagttctgctatgtggcgcggtattatcccgtattgacgccgggcaagagcaactcggtcgccgcatacactattctc
agaatgacttggttgagtactcaccagtcacagaaaagcatcttacggatggcatgacagtaagagaattatgcagtgctgccataaccatgagtgataacactgcggccaacttactt
ctgacaacgatcggaggaccgaaggagctaaccgcttttttgcacaacatgggggatcatgtaactcgccttgatcgttgggaaccggagctgaatgaagccataccaaacgacga
gcgtgacaccacgatgcctgtagcaatggcaacaacgttgcgcaaactattaactggcgaactacttactctagcttcccggcaacaattaatagactggatggaggcggataaagtt
gcaggaccacttctgcgctcggcccttccggctggctggtttattgctgataaatctggagccggtgagcgtgggtctcgcggtatcattgcagcactggggccagatggtaagccct
cccgtatcgtagttatctacacgacggggagtcaggcaactatggatgaacgaaatagacagatcgctgagataggtgcctcactgattaagcattggtaactgtcagaccaagttta
ctcatatatactttagattgatttaaaacttcatttttaatttaaaaggatctaggtgaagatcctttttgataatctcatgaccaaaatcccttaacgtgagttttcgttccactgagcgtcagac
cccgtagaaaagatcaaaggatcttcttgagatcctttttttctgcgcgtaatctgctgcttgcaaacaaaaaaaccaccgctaccagcggtggtttgtttgccggatcaagagctaccaa
ctctttttccgaaggtaactggcttcagcagagcgcagataccaaatactgttcttctagtgtagccgtagttaggccaccacttcaagaactctgtagcaccgcctacatacctcgctct
gctaatcctgttaccagtggctgctgccagtggcgataagtcgtgtcttaccgggttggactcaagacgatagttaccggataaggcgcagcggtcgggctgaacggggggttcgtg
cacacagcccagcttggagcgaacgacctacaccgaactgagatacctacagcgtgagctatgagaaagcgccacgcttcccgaagggagaaaggcggacaggtatccggtaa
gcggcagggtcggaacaggagagcgcacgagggagcttccagggggaaacgcctggtatctttatagtcctgtcgggtttcgccacctctgacttgagcgtcgatttttgtgatgctc
gtcaggggggcggagcctatggaaaaacgccagcaacgcggcctttttacggttcctggccttttgctggccttttgctcacatgttctttcctgcgttatcccctgattctgtggataacc
gtattaccgcctttgagtgagctgataccgctcgccgcagccgaacgaccgagcgcagcgagtcagtgagcgaggaagcggaagagcgcccaatacgcaaaccgcctctcccc
gcgcgttggccgattcattaatgcagctggcacgacaggtttcccgactggaaagcgggcagtga**GCGCAACGCAATTAATGTGAGTTAGCTCAC
TCATTAGGCACCCCAGGCTTTACACTTTATGCTTCCGGCTCGTATGTTGTGTGGAATTGTGAGCGGATAA
CAATTTCACACAGGAAACAGCTATGACCATGATTACGCCAAGCTTGCATGCCTGCAGGTCGACTCTAGA
GGATCCCCGGGTACCGAGCTCGAATTCACTGGCCGTCGTTTTACAACGTCGTGACTGGGAAAACCCTGG
CGTTACCCAACTTAATCGCCTTGCAGCACATCCCCCTTTCGC**cagctggcgtaatagcgaagaggcccgcaccgatcgcccttcccaa
cagttgcgcagcctgaatggcgaatggcgcctgatgcggtattttctccttacgcatctgtgcggtatttcacaccgcatatggtgcactctcagtacaatctgctctgatgccgcatagt
taagccagccccgacacccgccaacacccgctgacgcgccctgacgggcttgtctgctcccggcatccgcttacagacaagctgtgaccgtctccgggagctgcatgtgtcagag
gttttcaccgtcatcaccgaaacgcgcga

*upper cases + **boldened**: reference template for polymerase error rate measurement.

atto633_RP (17 bases):
/5ATTO633N/**ATTCACTGTCACGGCGC**

extension_template (22 bases):
TGATG**GCGCCGTGACAGTGAAT**

### Unique molecule indexed (UMI) primers for single-molecule polymerase error measurement:

| Name | Sequence | Purification | Length |
|---|---|---|---|
| **YX_F00** | ACACTCTTTCCCTACACGACGCTCTTCCGATCT**CTGT**NNNNNNNNNNNNNNNgcgcaacgcaattaatgtga | PAGE | 72 |
| **YX_F01** | ACACTCTTTCCCTACACGACGCTCTTCCGATCT**ACGTA**NNNNNNNNNNNNNNNgcgcaacgcaattaatgtga | PAGE | 73 |
| **YX_F02** | ACACTCTTTCCCTACACGACGCTCTTCCGATCT**ATTGAT**NNNNNNNNNNNNNNNgcgcaacgcaattaatgtga | PAGE | 74 |
| **YX_F03** | ACACTCTTTCCCTACACGACGCTCTTCCGATCT**AGACTGG**NNNNNNNNNNNNNNNgcgcaacgcaattaatgtga | PAGE | 75 |
| **YX_F04** | ACACTCTTTCCCTACACGACGCTCTTCCGATCT**CAGTCATC**NNNNNNNNNNNNNNNgcgcaacgcaattaatgtga | PAGE | 76 |
| **YX_F05** | ACACTCTTTCCCTACACGACGCTCTTCCGATCT**CGTCCACTG**NNNNNNNNNNNNNNNgcgcaacgcaattaatgtga | PAGE | 77 |
| **YX_F06** | ACACTCTTTCCCTACACGACGCTCTTCCGATCT**TACTGCATAC**NNNNNNNNNNNNNNNgcgcaacgcaattaatgtga | PAGE | 78 |
| **YX_F07** | ACACTCTTTCCCTACACGACGCTCTTCCGATCT**TCTACTGAGTA**NNNNNNNNNNNNNNNgcgcaacgcaattaatgtga | PAGE | 79 |
| **YX_F08** | ACACTCTTTCCCTACACGACGCTCTTCCGATCT**TGCAGCTCTAGT**NNNNNNNNNNNNNNNgcgcaacgcaattaatgtga | PAGE | 80 |
| **YX_F09** | ACACTCTTTCCCTACACGACGCTCTTCCGATCT**GTCCGGCAATCGG**NNNNNNNNNNNNNNNgcgcaacgcaattaatgtga | PAGE | 81 |
| **YX_F10** | ACACTCTTTCCCTACACGACGCTCTTCCGATCT**GAAGTCAGCGTACG**NNNNNNNNNNNNNNNgcgcaacgcaattaatgtga | PAGE | 82 |
| **YX_R00** | CGTGATGTGACTGGAGTTCAGACGTGTGCTCTTCCGATCT**GGTA**NNNNNNNNNNNNNNNgcgaaagggggatgtgc | PAGE | 76 |
| **YX_R01** | CGTGATGTGACTGGAGTTCAGACGTGTGCTCTTCCGATCT**TCAGT**NNNNNNNNNNNNNNNgcgaaagggggatgtgc | PAGE | 77 |
| **YX_R02** | CGTGATGTGACTGGAGTTCAGACGTGTGCTCTTCCGATCT**TGCCTG**NNNNNNNNNNNNNNNgcgaaagggggatgtgc | PAGE | 78 |
| **YX_R03** | CGTGATGTGACTGGAGTTCAGACGTGTGCTCTTCCGATCT**TAGAGCA**NNNNNNNNNNNNNNNgcgaaagggggatgtgc | PAGE | 79 |
| **YX_R04** | CGTGATGTGACTGGAGTTCAGACGTGTGCTCTTCCGATCT**CTAGCTGC**NNNNNNNNNNNNNNNgcgaaagggggatgtgc | PAGE | 80 |
| **YX_R05** | CGTGATGTGACTGGAGTTCAGACGTGTGCTCTTCCGATCT**CATCCTCGA**NNNNNNNNNNNNNNNgcgaaagggggatgtgc | PAGE | 81 |
| **YX_R06** | CGTGATGTGACTGGAGTTCAGACGTGTGCTCTTCCGATCT**GTCGACTGTC**NNNNNNNNNNNNNNNgcgaaagggggatgtgc | PAGE | 82 |
| **YX_R07** | CGTGATGTGACTGGAGTTCAGACGTGTGCTCTTCCGATCT**GCGTCGATAGT**NNNNNNNNNNNNNNNgcgaaagggggatgtgc | PAGE | 83 |
| **YX_R08** | CGTGATGTGACTGGAGTTCAGACGTGTGCTCTTCCGATCT**ATGCGACACTCA**NNNNNNNNNNNNNNNgcgaaagggggatgtgc | PAGE | 84 |
| **YX_R09** | CGTGATGTGACTGGAGTTCAGACGTGTGCTCTTCCGATCT**AGTACAGCTTAGA**NNNNNNNNNNNNNNNgcgaaagggggatgtgc | PAGE | 85 |
| **YX_R10** | CGTGATGTGACTGGAGTTCAGACGTGTGCTCTTCCGATCT**ACTAACTAGAGTCG**NNNNNNNNNNNNNNNgcgaaagggggatgtgc | PAGE | 86 |
