## Supplementary Files 1 - 8 for "Temperature Effect on Polymerase Fidelity": sfile_2_psychrophilic_protein_purification.pdf

Yuan Xue (Quake Lab - 2020/08/06)

**Note:** My protocol is adapted from the M9ZB protocol originally provided by Alison Ringel from Wolberger lab and HSP-removal protocol from Tony Hyman lab.

**Purpose:** To purify psychrophilic, large molecular weight protein with low solubility. I successfully purified a ~125 kDa MBP-fused psychrophilic polymerase which retained functional activity.

**Equipments:**

1. Shaking incubator for 10C and 37C
2. French press homogenizer system set up in cold room or with in-built refrigeration system.
3. Refrigerated liter volume centrifugation system such as Beckman Coulter JXN-26 centrifuge with JLA 8-1000 rotor
4. Refrigerated ultracentrifugation system (up to 235000 rcf) such as Beckman Coulter XL-100K Ultracentrifuge with Type 45 Ti, fixed angle rotor.
5. Tube rotator
6. **Preferred (optional):** Ultra yield baffled growth flask

**Materials:**

1. Solubility-tagged fusion construct (recommended vector: pMAL-c5X).
   **## This protocol assumes MBP-fusion tag is used. GST in my hand did not enhance solubility ##**
2. <u>Materials required by M9ZB protocol</u> (Alison Ringel, Wolberger lab)
   **## All solutions should be autoclaved or vacuum filtered prior to use ##**
3. Arctic Express Cell Line (DE3).
   **## Arctic Express cells carry a plasmid encoding for cold-adapted chaperons, thus enhancing protein stability at lower temperature range. This plasmid vector also carries a gentamycin selective marker ##**
   **## If protein contains rare codons, use Arctic Express Cell Lines optimized for rare codons (RIL and RP). ##**
4. Selective antibiotic for express vector
5. 20 mg/mL Gentamycin
6. 1M IPTG
7. Amylose resin

8. 1M Maltose solution
9. Dnase I (lyophilized powder is fine)
10. Halt protease inhibitor cocktail (Thermo fisher)
11. Adenosine 5′-triphosphate disodium salt hydrate
12. 40% glycerol, autoclaved
13. 1M Tris-Hcl, pH 7.50, filtered
14. 5M NaCl, filtered
15. 14.3M BME
16. 4M Kcl, filtered
17. miliQ water

**Procedure:**

Day 1

1. Heat shock transform Arctic Express (AE) cells with expression construct as per manufacturer’s instruction. Plate on LB agar with selective marker for your expression construct. **It is not necessary to add gentamycin to the LB plate**.
2. For each protein to be expressed, prepare 1 liter of autoclaved M9ZB media with only M9 salt, NaCl, and casamino acids. Set this aside and allow to cool on bench.

Day 2

1. In the morning, pick about 6 colonies from each transformed AE plate and inoculate each colony in 1 mL LB with selective antibiotic for expression and 20 ug/mL gentamycin. Incubate at 37C shaking at 250 rpm for 3-6 hours.
2. In the meantime, for each liter of M9ZB, prepare 50 mL MDG media without antibiotics added just yet.
3. After ~3 hours, it will be obvious that some AE cultures are not turbid (these likely lost their gentamycin plasmid during transformation). Pool the cultures that are turbid.
4. Transfer 100 uL of the LB pre-culture to 50 mL MDG. Add in selective antibiotic and 20 ug/mL gentamycin. Shake at 37C overnight.

Day 3

1. Add in glycerol, MgSO4, metal mix, and selective antibiotic for expression and 20 ug/mL gentamycin into M9ZB.
2. Transfer 10 mL of MDG culture into 1 liter of M9ZB. Shake at 37C for at least 4 hours until OD is >2.5.

**## In my hands, I have incubated M9ZB culture for at least 7 hours to reach OD 2.5 ##**

3. Once M9ZB has reached desired OD, place flask on ice for at least 30 minutes. Start cooling incubator down to 10C.
4. Collect 50 uL of culture as uninduced control for SDS-PAGE analysis. Heat on 95C for 5 mins in Laemelli buffer.
5. Once culture has chilled, add IPTG to the culture to 1 mM. Shake culture with 250 rpm at 10C for 24 hours.

Day 4

***All following steps should be conducted on ice or in cold room.**

1. Collect 50 uL of culture as induced control for SDS-PAGE analysis. Heat on 95C for 5 mins in Laemelli buffer.
2. Pellet M9ZB culture by centrifuging at 12 x krcf for 45 minutes at 4C (Beckman Coulter JXN-26, JLA 8-1000 rotor).
3. While centrifuging, make up 200 mL lysis buffer for one liter of M9ZB culture:

| | |
|---|---|
| 142 mL | cold miliQ water |
| 4 mL | 1M Tris-HCl, pH 7.50 |
| 50 mL | 40% Glycerol |
| 2 mL | 5M NaCl |
| 140 uL | 14.3M BME |
| ~400 U | DNase I |
| 2 mL | 100X protease inhibitor cocktail |

***Store in 4C until next step.**

4. Resuspend each liter of cell pellet in ~150 mL lysis buffer.
5. Lyse cells in French Press Homogenizer system. Pass cells through once at 500 psi, then two more times at 5000-10000 psi.
6. Collect 50 uL of lysate as total protein fraction for SDS-PAGE analysis. Heat on 95C for 5 mins in Laemelli buffer.
7. Clarify lysate by centrifuging at ~21 x krcf for 45 minutes at 4C (Beckman Coulter XL-100K Ultracentrifuge, Type 45 Ti rotor). Equilibrate amylose resin as per manufacturer's instructions.
8. Collect 50 uL of supernatant fraction and pick a "snob" of pellet fraction with pipette tip (dissolve in 50 uL Laemmeli buffer) for SDS-PAGE analysis. Heat on 95C for 5 mins in Laemelli buffer.
9. Bind supernatant fraction to 1 mL equilibrated amylose resin per liter of culture on tube rotator in cold room for 3 hours.

***It is important  to not bind lysate for overnight as some proteins can form aggregates**

10. Prepare 200 mL wash buffer:

| | |
|---|---|
| 144 mL | cold miliQ water |
| 4 mL | 1M Tris-HCl, pH 7.50 |
| 50 mL | 40% Glycerol |

| | |
|---|---|
| 2 mL | 5M NaCl |
| 140 uL | 14.3M BME |

Prepare 200 mL Cpn60-removal buffer:

| | |
|---|---|
| 135.5 mL | cold miliQ water |
| 10 mL | 1M Tris-HCl, pH 7.50 |
| 50 mL | 40% Glycerol |
| 2 mL | 5M NaCl |
| 2.5 mL | 4M KCl |
| 0.551 g | Adenosine 5′-triphosphate disodium salt hydrate (final: 5 mM) |
| 140 uL | 14.3M BME |

***Store both buffers in 4C until next step**

11. Pour resin mixture onto a gravity flow column. Collect flow-through on a separate flask.
12. Collect 50 uL of flow-through fraction for SDS-PAGE analysis. Heat on 95C for 5 mins in Laemelli buffer.
13. Wash amylose resin with 10 column volume of wash buffer. Collect 50 uL of wash fraction #1 and 50 uL of amylose resin for SDS-PAGE analysis. Heat on 95C for 5 mins in Laemelli buffer.

**Often times, the chaperon (Cpn60) expressed in AE cell line gets co-purified with protein of interest. If removal of chaperon is not necessary, steps 14 and 15 can be skipped.**

####

14. Wash amylose resin with 20 column volume of Cpn60-removal buffer. Collect 50 uL of Cpn60-removal fraction for SDS-PAGE analysis. Heat on 95C for 5 mins in Laemelli buffer.
15. Wash amylose resin again with 10 column volume of wash buffer. Collect 50 uL of wash removal fraction #2 for SDS-PAGE analysis. Heat on 95C for 5 mins in Laemelli buffer.

####

16. Resuspend resin mixture in 5 mL wash buffer. Transfer to a clean conical tube.
17. Add 50 uL 1M maltose (final: 10 mM) to resin mixture and rotate in cold room for 3 hours.
18. Collect elution fraction in a new tube. Collect 50 uL of elution fraction for SDS-PAGE analysis. Heat on 95C for 5 mins in Laemelli buffer.

***Eluted fraction can be further purified in FPLC system with anion exchange columns and/or gel-filtration columns.**

19. Analyze protein fractions in SDS-PAGE gel. In resin fraction, there is generally additional product at ~60 kDa (Cpn60) and ~43 kDa (MBP). If Cpn60 removal step was complete, the 60 kDa band should be absent in the final elution fraction. Presence of 43 kDa (MBP) indicates that protease inhibition was not complete. This can generally be removed by gel-filtration or Amicon filter unit.

20. At this step, I generally dialyze against wash buffer and concentrate in Amicon centrifugal filter unit to desired concentration. Proteins can be snap-frozen in liquid nitrogen and stored in -80C.
